# The Host-specific Microbiota is Required for Diet-Specific Metabolic Homeostasis

**DOI:** 10.1101/2023.11.05.565654

**Authors:** Na Fei, Bingqing Xie, Tyler J. Long, Marissa StGeorge, Alan Tan, Sumeed Manzoor, Ashley M. Sidebottom, Melanie Spedale, Betty R Theriault, Dinanath Sulakhe, Eugene B. Chang

## Abstract

In complex mammals, the importance and host-specificity of microbial communities have been demonstrated through their positive effects on host immune fitness or performance. However, whether host metabolic physiology homeostasis depends on a specific bacterial community exclusive to the host remains unclear. Here, we show that the coevolved host-specific microbiota is required to maintain diet-specific flexible and sufficient metabolic homeostasis through a high colonization rate, modulating gut metabolites, and related targets. Using germ-free (GF) mice, we tested whether the fitness benefiting the host metabolic phenotype of microbiota was host-specific. We demonstrated that GF mice associated with exogenous microbiota (human microbiota (HM)), which exhibited different and reduced gut microbial species diversity, significantly elevated metabolic rate, and exhibited metabolic insufficiency, all characteristics of GF mice. Strikingly, the absence of the host-specific microbiome attenuated high-fat diet-specific metabolism features. Different diets caused different metabolic changes in only host-specific microbiota-associated mice, not the host-microbiota mismatched mice. While RNA sequencing revealed subtle changes in the expression of genes in the liver, GF mice and HM mice showed considerably altered expression of genes associated with metabolic physiology compared to GF mice associated with host-specific microbiota. The effect of diet outweighed microbiota in the liver transcriptome. These changes occurred in the setting of decreased luminal short-chain fatty acids (SCFAs) and the secondary bile acid (BAs) pool and downstream gut signaling targets in HM and GF mice, which affects whole-body metabolism. These data indicate that a foreign microbial community provides little metabolic benefit to the host when compared to a host-specific microbiome, due to the colonization selection pressure and microbiota-derived metabolites dysfunction. Overall, microbiome fitness effects on the host metabolic phenotype were host-specific. Understanding the impact of the host-specificity of the microbiome on metabolic homeostasis may provide important insights for building a better probiotic.

**Highlights:**

- Microbiome fitness effects on the host metabolic phenotype were host-specific in mammals.
- Human microbiota-associated mice exhibited lower host metabolic fitness or performance, and similar functional costs in GF mice.
- Different diets cause different metabolic changes only in host-specific microbiota-associated mice, not the host-microbiota mismatched mice.
- The defective gut microbiota in host-specific microbiota, microbial metabolites and related targets likely drive the metabolic homeostasis.

**Graphical Abstract:** 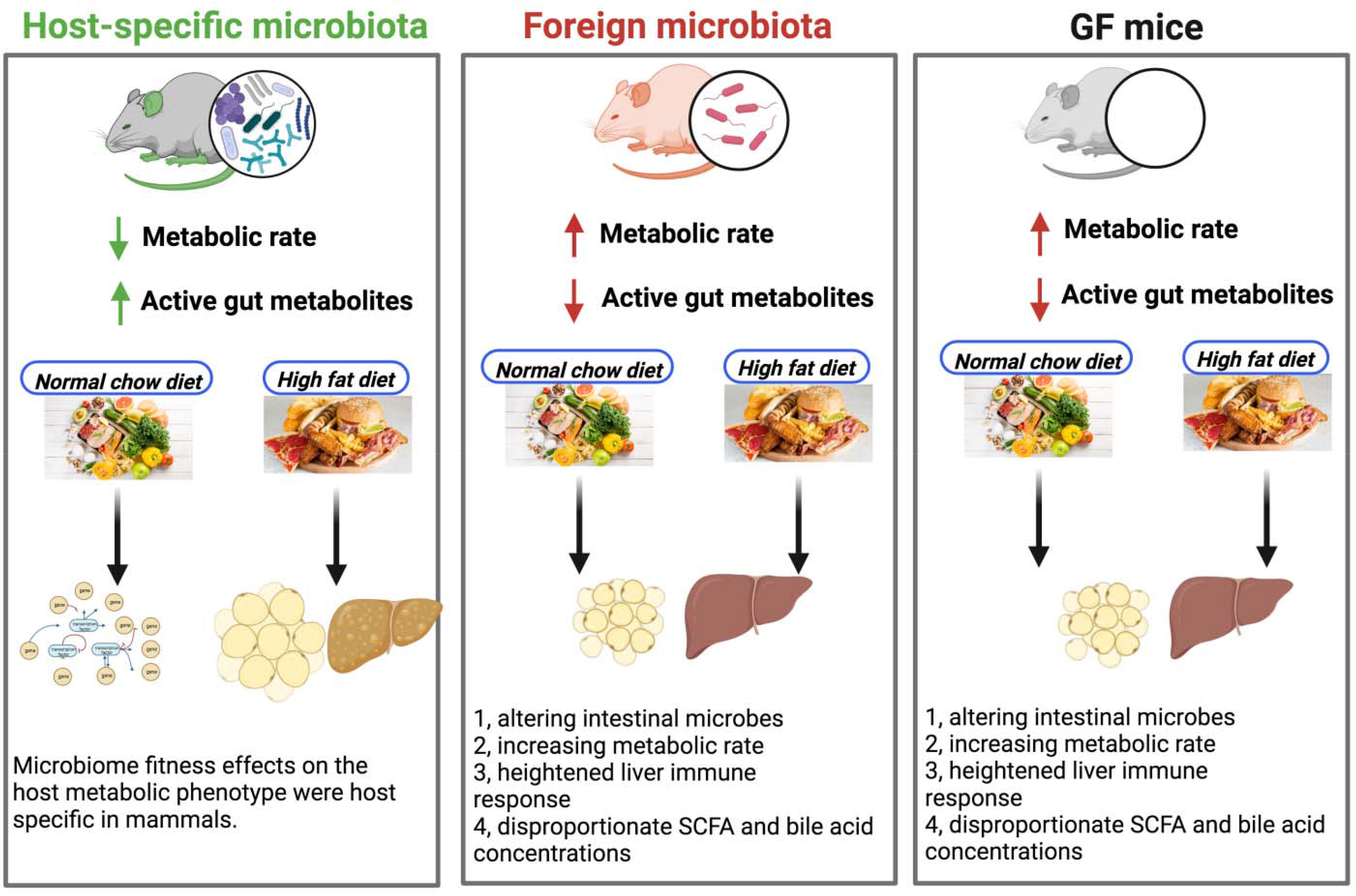

## Introduction

Phylosymbiosis relationships between the host and the microbial community habited in their gut have been documented in mammals ^1–3^. In mammals, the vast microbes in their gut may govern important and positive functions critical to the host’s health status, including development, immunity, and nutrition ^4^. It is becoming increasingly evident that the gut microbiome contributes specifically to the host’s performance and fitness. For example, the host-specific gut microbial community has been documented to positively affect their mammal hosts by altering host immune system maturation ^5^. However, it is unclear whether the community typically colonizing a given mammalian host species preferentially stimulates a specific program of host metabolic maturation.

Accumulating evidence has shown the congruent relationships between hosts’ metabolism and associated microbial communities ^6^. Host gut ecology and the nutritional composition of the diet determined the variations of the commensal host-specific intestinal microbiota composition and metabolic activity ^7^. Indeed, germ-free (GF) or antibiotic-treated mice displayed developmental defects in abnormal nutrient absorption and metabolic deficiencies in the host and thus exhibited resistance to high-fat diet-induced obesity relative to the mice colonized with a health-associated mouse commensal microbiota ^8–10^. Gene expression patterns in metabolic physiology in the liver and gut of GF or antibiotic-treated mice are distinct from conventionally raised mice ^8–10^. However, the effects of the host-microbiome mismatch on the host metabolism have not been explored. It remains unclear whether and to what extent the health-associated metabolic development depends on the specific bacterial community exclusive to the host. In the present study, we aimed to investigate if mammalian metabolic homeostasis is dependent on the presence of a coevolved host-specific microbiota uniquely capable of stimulating metabolic signaling, to provide further insights into the fitness effects of host-specific microbiomes. With a better understanding of the effects of host-specific microbiota on host metabolic physiology, mice associated with exogenous microbiome can be more effectively used in animal experiments and provide important insights for building a better probiotic.

Gut microbial metabolites play a critical role in their host metabolic physiology homeostasis ^11^. GF or antibiotic-treated mice displayed microbial dysmetabolism compared with conventionally colonized mice, including decreased luminal and serum short-chain fatty acid (SCFA) and secondary bile acid (BA) pool ^12,13^. Those gut metabolite changes mechanistically lead to altered glucose and metabolic homeostasis in the host ^12,13^. First, gut microbes are responsible for additional calorie extraction through the fermentation of indigestible carbohydrates into SCFAs, which can serve as essential fuel for the colonic cells, through the SCFA receptors, G protein-coupled receptors (GPR41, GPR43, and GPR109A) ^14^. Second, the microbiota can also affect metabolic homeostasis by altering the BA pool ^15^. Intestinal microbiota can deconjugate BAs and convert primary BAs into secondary ones, which are known mediators of glucose metabolism ^15^. BAs can affect host metabolism by signaling through the farnesoid X receptor (FXR) and G-protein-coupled BA receptor (TGR5) ^15^. Thus, it is critical to investigate if the host-microbiota mismatched mice have a luminal dysmetabolism state similar to what is described in GF mice, hence, applying markedly influence on metabolic regulation.

Earlier mouse studies showed that in the case of diet-induced obesity, GF mice could protect against dysmetabolism under the same diet condition ^8^. However, exogenous microbiota-associated mice differ from GF mice in two ways: (a) exogenous microbiota-associated mice do not have complete depletion of their gut microbiome, and (b) exogenous microbiota themselves have a direct effect on host metabolic homeostasis. If probiotics, particular bacterial species targeting, or fecal microbial transplantation are therapeutic approaches for diseases, principles are required for the host-related microbiome to reconstitute the microbiota with those exogenous particular species. Thus, knowing the physiological effects of exogenous microbiota on normal metabolism is a prerequisite for implementing these potential therapies.

In this study, we aim to investigate the fitness benefits of the host-specific microbiota on host metabolism homeostasis by introducing human (HM) or mouse microbiota (MM) in normal-chow or high-fat-fed GF mice, with GF mice (GF) as control. We used metabolic cage, liver transcriptome (RNA-seq), and gut metabolome to compare the microbiota’s effect on the host metabolic status of MM mice, HM mice, and GF mice. Our results showed that microbiome fitness effects on the host metabolic phenotype were host-specific. We found that the metabolic rate was globally affected, and hepatic genes involved in crucial processes exhibited an altered expression profile in HM male mice. Strikingly, we observed a similar metabolic status between HM male mice and GF male mice and confirmed an attenuation of metabolic activities under the high-fat diet in these mice. Furthermore, HM-associated mice consume a significant microbiota depletion. These changes in metabolic homeostasis are associated with altered gut luminal BA and SCFA profiles. There was a consequence of altered metabolic signaling, likely due to the defective gut microbial maturation of HM and GF mice. Last, we observed different diets that affected the host specificity of microbial only on host metabolic physiology in different ways. These observations suggest that mammalian hosts have coevolved with a specific consortium of bacterial species uniquely capable of stimulating metabolic homeostasis.

## Results

### Host-microbiota mismatch feature plays a role in the development of host metabolic impairment

To determine the influence of host-specific microbial colonization on host metabolic homeostasis, germ-free (GF) C57Bl/6J male and female mice underwent oral gavage with pooled cecal specimens from either healthy humans (HM) or specific pathogen-free (SPF) C57Bl/6J mice (MM), with GF mice as control. The recipient mice were then maintained in separate gnotobiotic isolators under normal chow diet (NCD) or high-fat diet (HFD) treatment for 8 weeks (**Figure 1A**). Metabolic parameters were evaluated to determine the consequence of different microbiota on host metabolic status. We did not observe significant differences in body weight, liver weight, or adipose tissue weight between NCD-fed male mice harboring human or mouse microbiota compared to GF male mice (p>0.05, **Figure 1B to H**). However, significant differences in body weight were observed among HFD-fed male mouse groups; HFD-fed-MM male mice exhibited significantly higher body weight compared to HFD-fed HM or GF male mice (p<0.05, **Figure 1B**). MM male mice on HFD had gained more than 6% weight compared to HFD-fed HM male mice and around 10% compared to their GF counterparts. Furthermore, no significant difference in body weight was observed between HFD-fed HM and GF male mice (p>0.05, **Figure 1B**), showing that the exogenous microbial community (HM) does not have as detrimental effects as the indigenous one on the body weight even under HFD treatment in male mice. In parallel, HFD-fed MM male mice showed elevated liver weight compared to HFD-fed HM or GF controls (**Figure 1C**). HFD-fed HM male mice displayed higher liver weight compared to HFD-fed GF controls; however, still significantly lower than HFD-fed MM male mice. Similarly, adipose tissue (retroperitoneal, gonadal, inguinal, and mesenteric fat) exhibited similar weight among GF, HM, and MM male mouse groups under NCD-fed; however, significant increases in retroperitoneal and inguinal fat were observed in HFD-fed MM male mice compared to HFD-fed HM or GF male mice (p<0.05, **Figure 1D-G**). Total adipose tissue showed similar results. A significant increase in the total adipose depot was observed in HFD-fed MM male mice compared to the GF counterparts (p<0.05, **Figure 1H**), but there were no differences between HFD-fed HM and GF male mice (p>0.05, **Figure 1H**).

**Fig. 1:**
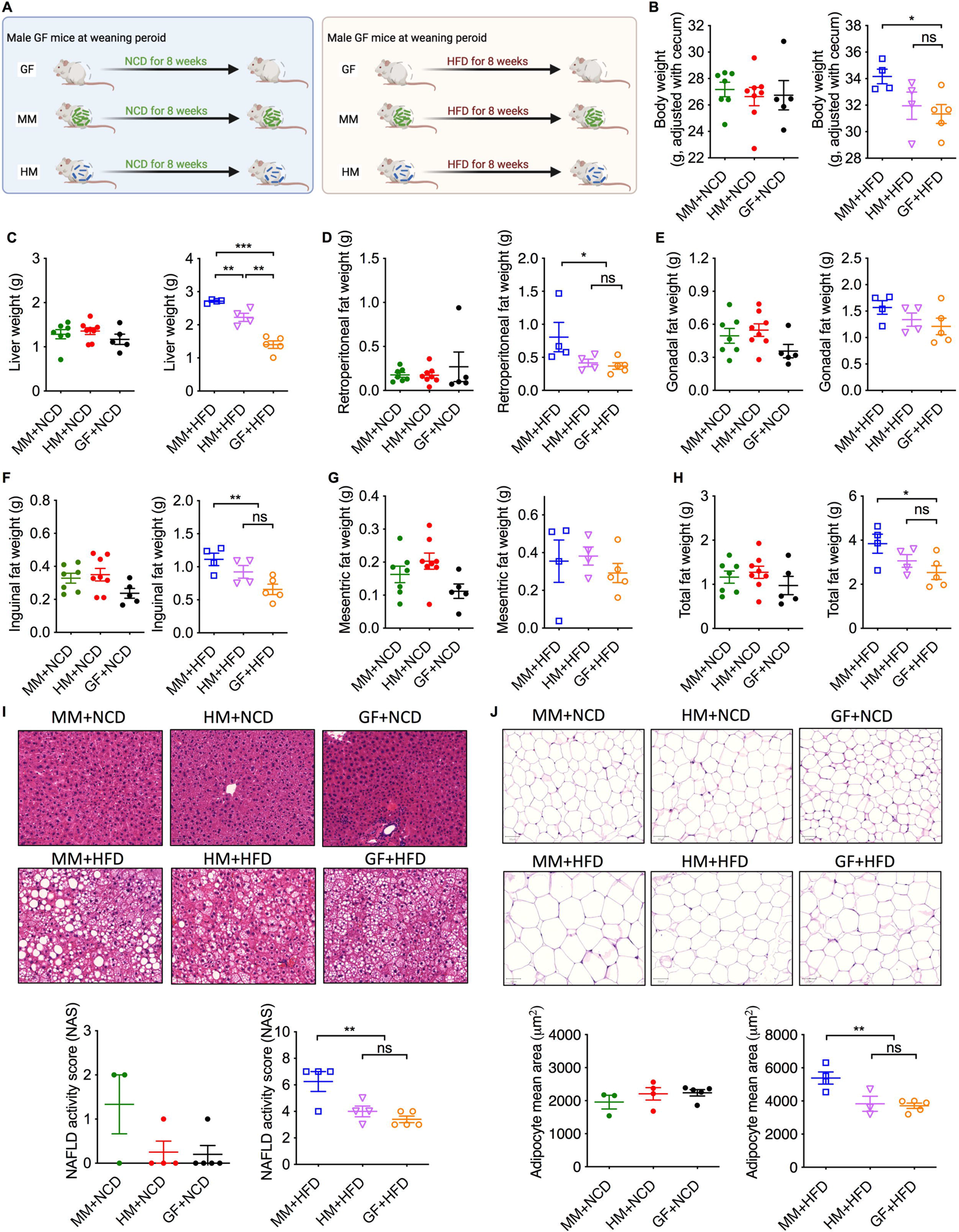
The host-specific microbiota is required for diet-specific metabolic syndrome in male mice. **A.** Experimental scheme. Wild-type GF male mice were given either human, mouse cecal contents, or PBS at 3 weeks of age and treated with a normal chow diet (NCD) or high-fat diet (HFD) for 8 weeks. **B**. Body weight of NCD-fed or HFD-fed HM, MM, and GF mice after 8 weeks with a diet treatment. **C**. Liver weight of NCD-fed or HFD-fed HM, MM, and GF male mice. **D-H**. Adipose tissue weight, retroperitoneal fat (**D**), gonadal fat (**E**), inguinal fat (**F**), mesenteric fat (**G**), and total fat (**H**) of NCD-fed or HFD-fed HM, MM, and GF male mice. I. Formalin-fixed liver hematoxylin and eosin stain (H&E, up) and corresponding graphs representing NAFLD activity score (NAS, down) of NCD-fed or HFD-fed HM, MM, and GF male mice. Scale bar, 100 μm. (Representative images are shown for livers from each group. **J**. Formalin-fixed gonadal fat hematoxylin and eosin stain (H&E, up) and corresponding graphs representing NAFLD activity score (NAS, down) of NCD-fed or HFD-fed HM, MM, and GF male mice. Scale bar, 100 μm. (Representative images are shown for Gonadal fats from each group. All microscopy images were at 100x magnification). All microscopy images were at 100x magnification, Scalebars = 100 μm. Results are expressed as the mean ± SEM. n = 4–6 male mice per group. Statistical comparison was performed by testing normality using Kolmogorov–Smirnov test and then ANOVA or Kruskal–Wallis test with Turkey’s post hoc test. *p < 0.05. *p ≤ 0.05, **p ≤ 0.01, and ***p ≤ 0.001. ns, not significant. Gray shade represents 12-hour dark phases.

In the host-specific microbiota-associated male mice, liver histology was altered, characterized by a significantly higher NAFLD activity score in HFD-fed MM mice compared to the same diet-fed HM or GF mice (p<0.05, **Figure 1I**). Host-specific microbial mismatch features did not significantly dampen hepatic steatosis under HFD (**Figure 1I**). In terms of adipocyte effect, HFD-treated HM or GF male mice showed significantly lowered adipocyte size compared to MM mice, underscoring the importance of host-specific microbiota in promoting obesity (p<0.05, **Figure 1J**). We did not observe significant differences in the liver or adipocyte histology between NCD-fed male mice harboring human or mouse microbiota compared to GF male mice (p>0.05, **Figure 1I to J**). Regardless of metabolic change in male mice, however, the host-mismatch microbial feature did not affect metabolic homeostasis in HFD-fed female mice (**Figure S1**), which indicates the sex-specific phenomenon. The host-specific microbiota-associated mice, both male and female, exhibited regular cecum size and colon length (**Figure S2**). However, HM and GF male mice displayed enlarged cecum size and longer colon length than their MM littermates (**Figure S2**). These results indicate that the host-specific microbiota is required for cecum and colon morphology in mice. Altogether, these results indicated that host-specific microbiota potentiated HFD-induced host metabolic dysfunction, with significant dysregulation of metabolic homeostasis and liver function. The host-specific microbial mismatch feature partly masked HFD’s adverse metabolic effects on the host in male mice.

### Host-microbiota mismatch feature results in higher energy expenditure and significant metabolic insufficiency and inflexibility

We next assessed the metabolic rate of these mice associated with different microbiota by placing them in metabolic chambers (Promethion SABLE systems) at ambient temperature for 24 hours. We demonstrated that, under HFD treatment, host-specific microbiota-associated male mice (MM) had reduced O_2_ consumption (VO_2_), CO_2_ production (VCO_2_) in both light and dark cycles, and energy expenditure during the night (**Figures 2B, D, and F**). No differences were observed between HM and GF male mice for these metabolic parameters under either NCD or HFD treatment (**Figures 2A-F**). A slight increase in VO_2_ and energy expenditure was noted in the HM male group relative to those in the GF group; however, no significant difference was observed (**Figures 2B, D, and F**). The lack of host-specific microbiota in HM and GF mice demonstrated that the exogenous or missing microbiota produced a substantial increase in metabolic rate in terms of enhanced VO2 and energy expenditure. That being said, all mice displayed similar levels of CO_2_ production (**Figures 2A**), VO_2_ (**Figure 2C**), and energy expenditure (**Figure 2E**) between the light and dark cycles under NCD treatment. Next, we have collected data on the respiratory exchange ratio (RER) among different microbiota-associated mice and GF mice to evaluate the differential level of nutrient substrate oxidation (which energy source among carbohydrates, lipids, or proteins is predominantly oxidized). Host-specific microbiota-associated male mice (MM) displayed a higher RER compared to HM and GF male mice in the light cycle under NCD treatment and across the entire light-dark cycle under HFD treatment, suggesting increased utilization of lipid as an energy source over carbohydrates in HM and GF male mice compared with that in MM male mice (Figure 2G-H). Exogenous microbiota (HM) caused a reduction in RER that was similar in GF mice in HFD treatment. Because the RER diminished and more O_2_ was consumed in HM and GF mice, our results indicated a decrease in carbohydrate fermentation, an increase in fatty acid oxidation, and promoted the use of lipid storage as an energy source in a lack of host-specific microbiota. Moreover, the enhancement of energy expenditure in HM and GF mice is not attributable to significant changes in food intake (expressed as kcal/day/mouse, **Figure 2K-L**). Because metabolic rate is associated with motor behavior, we further investigated whether physical activity differed among various groups. There was no significant difference in pedspeed among microbiotas in either the light or dark cycle under NCD (**Figure 2I**). Pedometer data showed a significant enhancement at nighttime in the HM and GF groups compared to that in the MM groups under HFD (**Figure 2J**). Similar to metabolic phenotype change, the host-mismatch microbial feature did not affect energy expenditure in female mice either (**Figure S3**). These experiments underlined three major metabolic traits in host-microbiota mismatched mice exhibiting a similar phenotype to GF mice: hypermetabolism, metabolic insufficiency, and inflexibility.

**Fig. 2:**
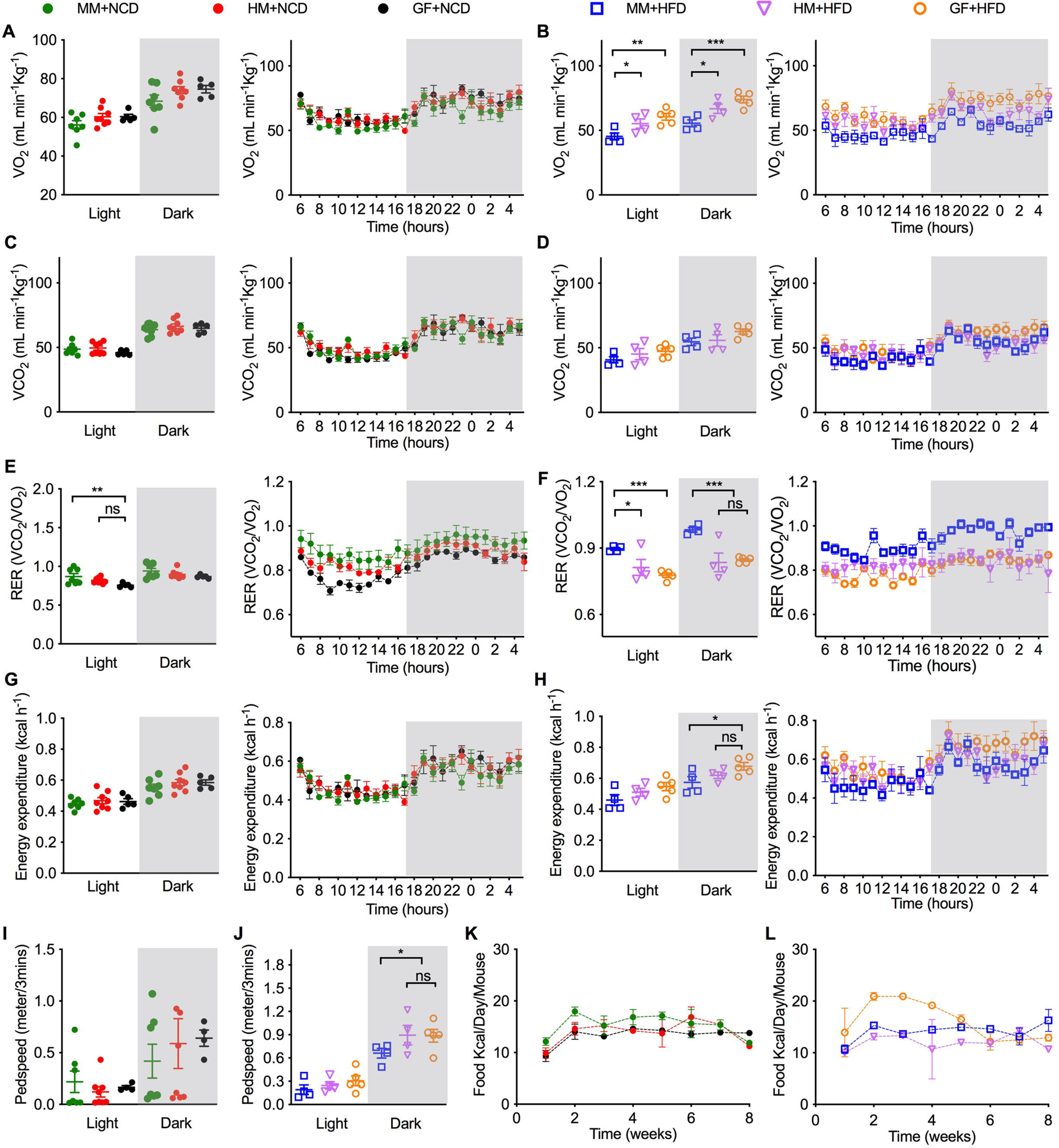
Mice lacking the matched host-specific microbiota displayed increased energy expenditure in HM male mice on a high-fat diet (HFD) compared to MM male mice. **A-B**. Mean O_2_ consumption (VO_2_) during light and dark phases and VO_2_ curves representing 24 consecutive hours of NCD-fed (**A**) or HFD-fed male mice (**B**). **C-D**. Mean VCO_2_ during light and dark phases and VCO_2_ curves representing 24 consecutive hours of NCD-fed (**C**) or HFD-fed male mice (**D**). **E-F**. Mean respiratory exchange ratio (RER = VCO_2_/VO_2_, where VCO_2_ is CO_2_ production) during light and dark phases and curves representing 24 consecutive hours of NCD-fed (**E**) or HFD-fed male mice (**F**). **G-H**. Mean energy expenditure during light and dark phases and curves representing 24 consecutive hours of NCD-fed (**G**) or HFD-fed male mice (**H**). **I-J**. Spontaneous total activity (meter/3mins) during light and dark phases of NCD-fed (**I**) or HFD-fed male mice (**J**). K-L. Food intake of NCD-fed (K) or HFD-fed male mice (**L**). Results are expressed as the mean ± SEM. n = 4–6 male mice per group. Statistical comparison was performed by testing normality using Kolmogorov–Smirnov test and then ANOVA or Kruskal–Wallis test with Turkey’s post hoc test. *p < 0.05. *p ≤ 0.05, **p ≤ 0.01, and ***p ≤ 0.001. ns, not significant. Gray shade represents 12-hour dark phases.

### The host-specific microbiota exhibit significantly different features and colonization rate than the exogenous microbiota

To evaluate the differences between the host-specific and non-specific composition of the gut microbiota and the contribution of diet to changes after being introduced to GF male recipients, we investigated the microbial structure using the 16S rRNA gene sequencing. Shannon diversity (a measure of α-diversity) assessment of two different groups of microbiomes showed that the MM mice had the greater diversity, with a significant decrease in the HM mice under NCD or HFD treatment (**Figures 3A**). The principal coordinate analysis of another measure of diversity, UniFrac distances, showed that the microbiomes of HM and MM-associated mice were quite distinct (**Figures 3B and C, left panel**). Pairwise comparison based on permutation tests (permutational multivariate analysis of variance (PERMANOVA)) showed significantly different weighted UniFrac distances between HM sand MM mouse samples under NCD or HFD treatment (**Figures 3B and C, right panel**). The relative abundance at the genus level was considerably different between MM and HM mice (**Figure 3D to G**). We observed that the host-specific microbiota-associated mice (MM) were dominated by *Bacteroides* and *Akkermansia* species under NCD or HFD treatment. In contrast, unclassified genera belonging to *S24-7*, *Clostridiales*, and *Oscillospira* were dominated in the HM mice under NCD or HFD treatment (**Figure 3D to G**). Both MM and HM mice microbiomes had a compositional shift (**Figure 3D and E**) with NCD or HFD treatment. These results indicate that exogenous microbiota-associated mice consume a significant microbiota depletion, especially of Firmicutes and Bacteroidetes species, decreasing microbial diversity of the microbiota.

**Fig. 3:**
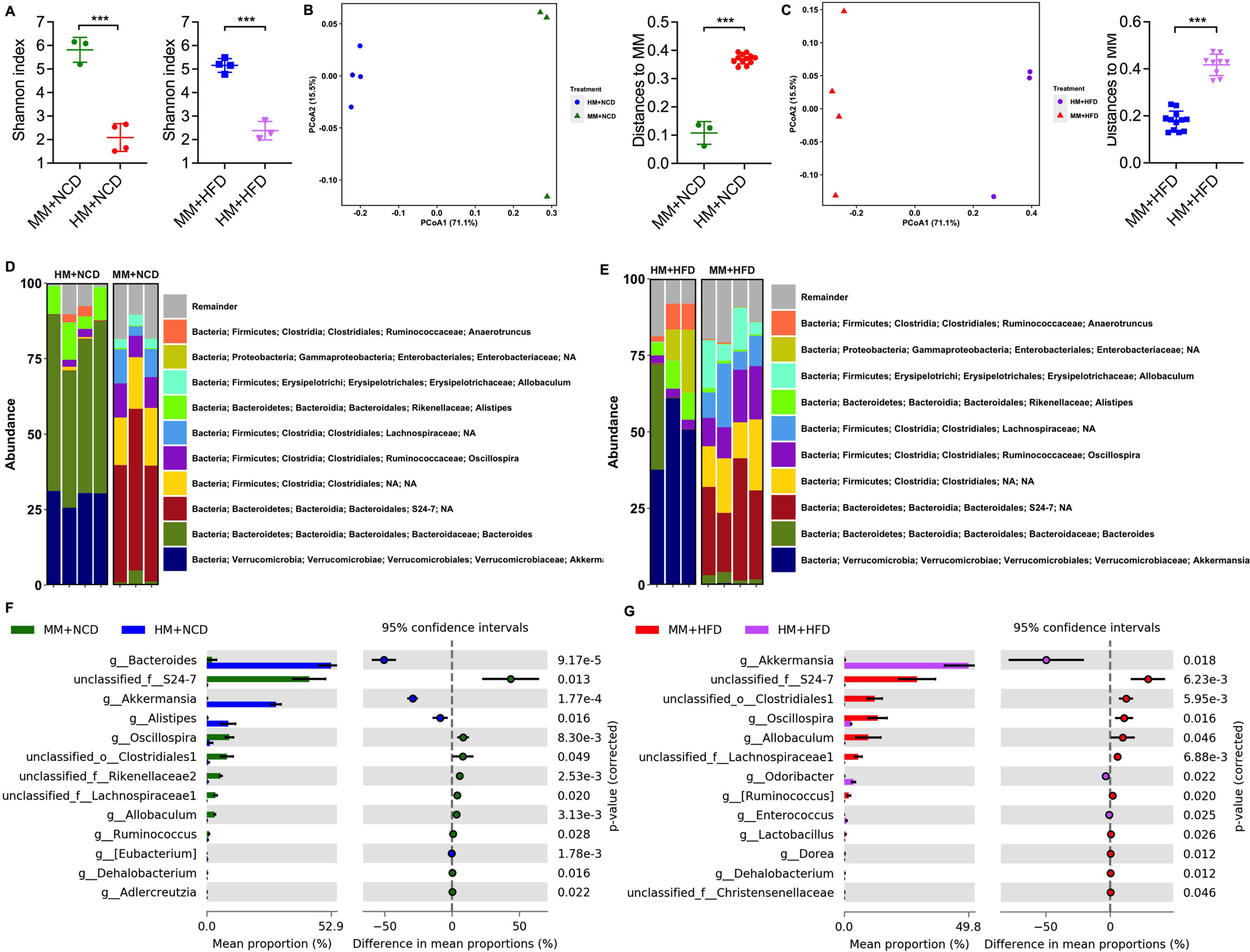
Microbiota composition distinct significantly between MM and HM. **A**. Cecal microbiota alpha diversity (Shannon index) of NCD-fed or HFD-fed HM and MM mice. **B-C**. PCoA plot (Weighted UniFrac distance) of cecal microbiota of NCD-fed (**B**) or HFD-fed HM and MM mice (**C**). **D-E**. Bar graph of bacterial abundance at genus level of NCD-fed (**D**) or HFD-fed HM and MM mice (**E**). **F-G**. Bacterial taxa differentially enriched in NCD-fed (**F**) or HFD-fed (**G**) HM compared to MM in mice. Results are expressed as the mean ± SEM. n = 3–4 male mice per group. Statistical comparison was performed by testing normality using Kolmogorov–Smirnov test and then ANOVA or Kruskal–Wallis test with Turkey’s post hoc test. *p < 0.05. *p ≤ 0.05, **p ≤ 0.01, and ***p ≤ 0.001. ns, not significant. Gray shade represents 12-hour dark phases.

### Exogenous gut microbiota induces significant changes in the transcriptome of the liver

To further explore how host-specific gut microbiota and diet influence the transcriptome in the liver in mice, we performed liver transcriptome profiling with RNA-seq. The multidimensional scaling analysis (MDS) demonstrated that diet’s effect was much more significant compared to various microbiotas on liver transcriptome (**Figure 4A**). Regarding the impact of various microbiotas, we found a clear separation between the MM, HM, and GF mice on NCD and a similar pattern between these mice on HFD (**Figure 4B**). Significant gene expression differences between HM, MM, and GF mice on different diets were identified using log2 fold change. Hierarchical clustering of all the DEGs and pairwise volcano plots revealed markedly distinct hepatic gene expression patterns in the MM group on NCD compared to those of HM and GF groups (**Figure 4C**), confirming major changes in hepatic gene expression induced by exogenous or missing gut microbiota. HM mice treatment with the NCD changed their expression towards levels seen in GF mouse livers. However, under HFD treatment, only a few differences were observed in the liver transcriptome between mice harboring human or mouse microbiota compared to GF male mice, which further underscored the effect of high-fat diet on the liver transcriptome outweighed that of different microbiotas. The volcano plot reports the upregulated and downregulated DE RNAs induced by exogenous gut microbiota and HFD in the liver of gnotobiotic mice (**Figure 4D**). 1816 (MM+NCD vs. HM+NCD (FDR p<0.05)) genes were dysregulated, of which 590 were downregulated in the HM group of mice, and 1226 were upregulated in the HM group (**Figure 4D**). In other comparisons, 181 (MM_NCD vs. GF_NCD), 14 (HM_HFD vs. GF_HFD), and 2 (MM_HFD vs. GF_HFD) genes were dysregulated (**Figure 4D**). We found no dysregulated genes induced by HM under HFD compared to the MM control group (**Figure 4D, Figure 5A** and **B**). Cross-comparison of the lists of genes differentially expressed in MM, HM vs. GF mice revealed 134 genes common in these two comparisons (MM_NCD vs. HM_NCD) and DEGs (MM_NCD vs. GF_NCD), with 1729 genes unshared to the 2 data sets (**Figure 4D, Figure 5A** and **B**).

**Fig. 4:**
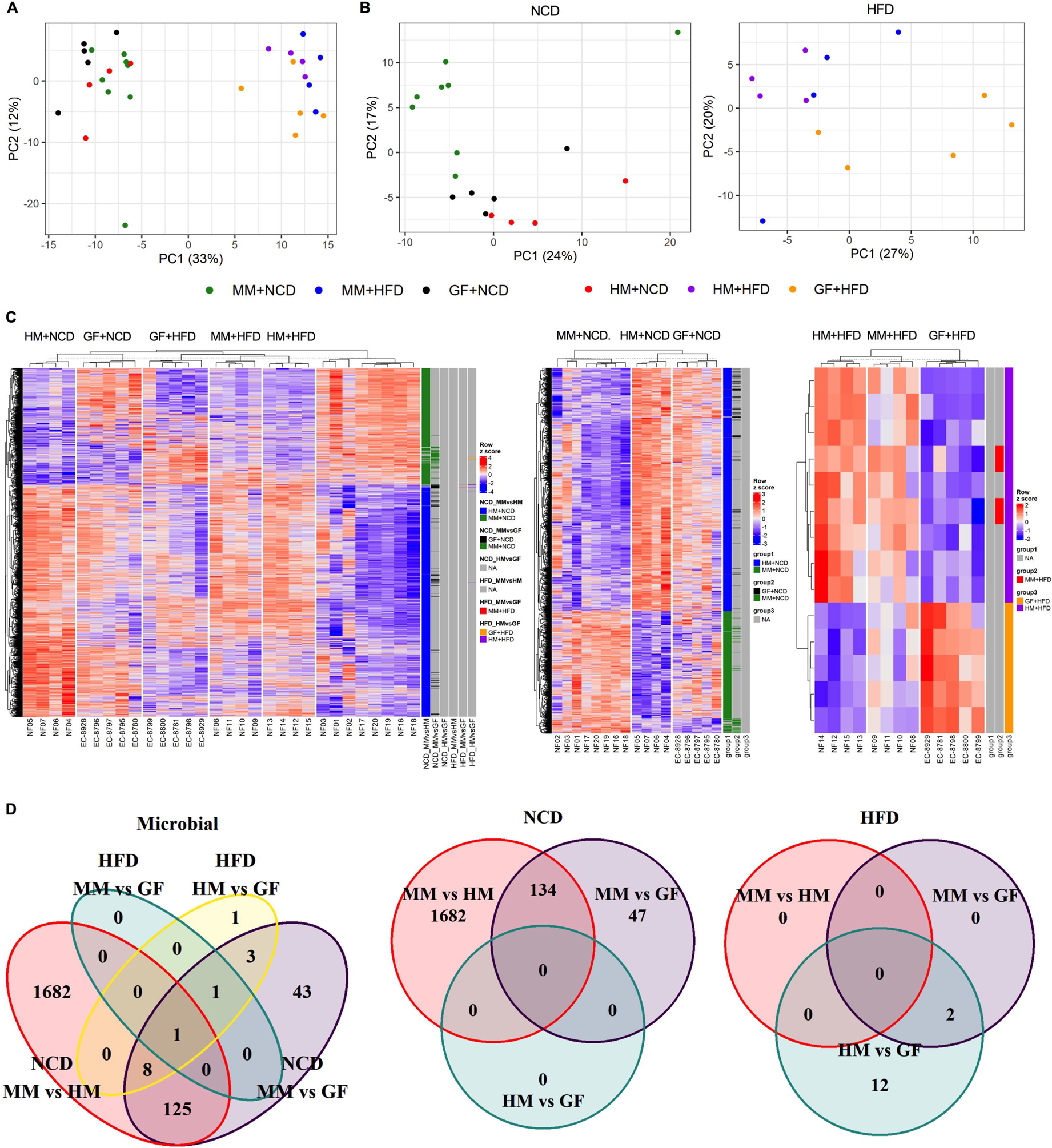
Non-matched microbiota induces major changes in the transcriptome of the liver. **A-B**. MDS analysis of the liver RNA-sequencing data of NCD-fed (**A**) or HFD-fed (**B**) HM, MM, and GF mice. **C**. Heatmap of differentially expressed genes in the liver of NCD-fed or HFD-fed HM, MM, and GF mice, generated using unsupervised hierarchical clustering. **D**. Venn diagram indicating the number of significant differentially expressed genes and the overlap between each set of genes of NCD-fed or HFD-fed HM, MM, and GF mice. **E**. Volcano plot showing the upregulated and downregulated DE RNAs of NCD-fed or HFD-fed HM, MM, and GF mice. **F**. Functional annotation of differentially expressed genes in the liver of NCD-fed or HFD-fed HM, MM, and GF mice identified by KEGG pathways. The top 10 KEGG functions are shown. KEGG, Kyoto Encyclopedia of Genes and Genomes. Adjusted p-value (corrected for multiple hypothesis testing with the Benjamin–Hochberg method) < 0.05. Count: number of differentially expressed genes within a given KEGG pathway; Grey dots within plot indicate insignificant (FDR p-value cutoff = 0.05) enrichment of KEGG pathway.

**Fig. 5:**
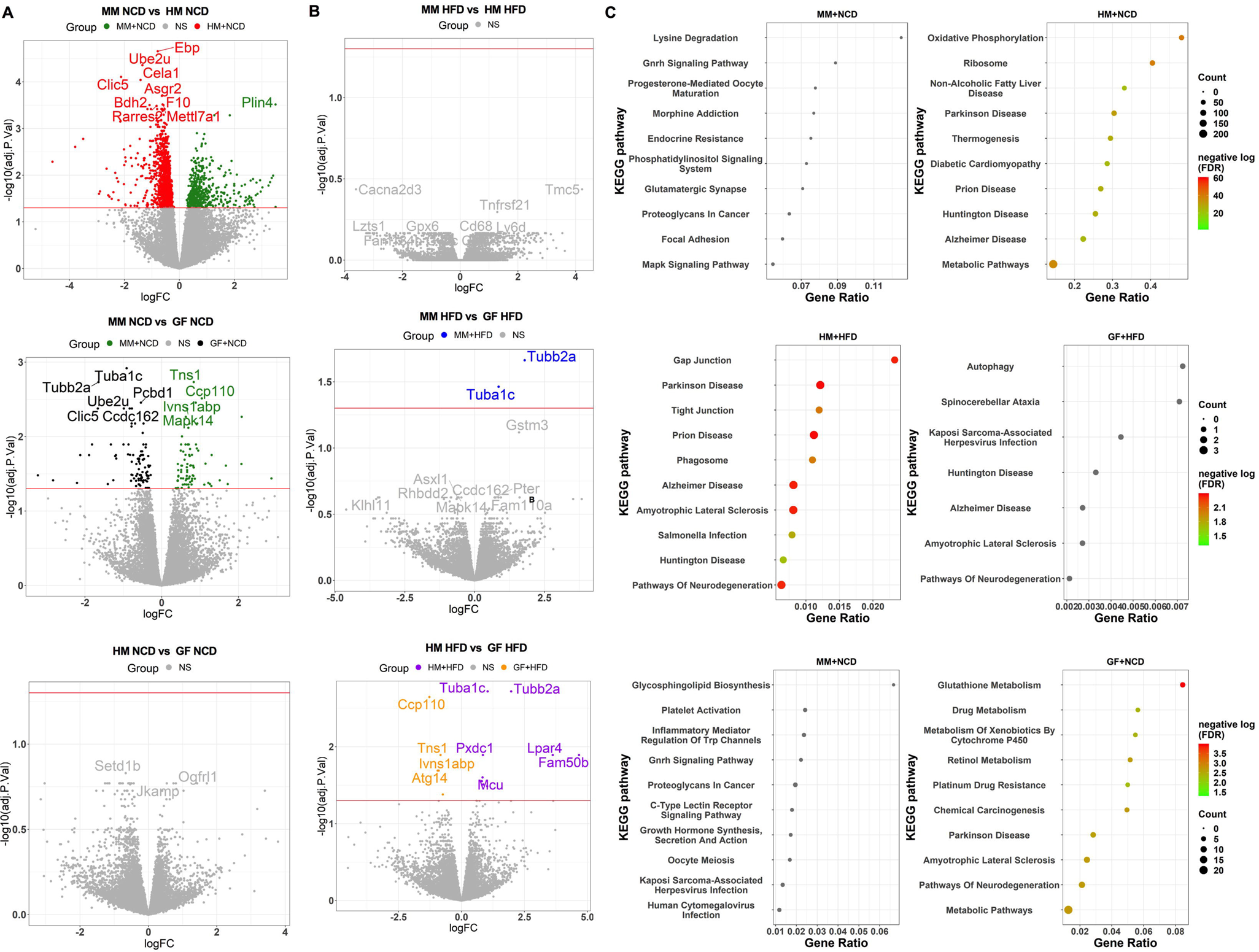
Non-matched microbiota induces metabolic dysfunction in the liver. **A** and **B**. Volcano plot showing the upregulated and downregulated DE RNAs of NCD-fed (**A**) or HFD-fed (**B**) HM, MM, and GF mice. **C**. Functional annotation of differentially expressed genes in the liver of NCD-fed or HFD-fed HM, MM, and GF mice identified by KEGG pathways. The top 10 KEGG functions are shown. KEGG, Kyoto Encyclopedia of Genes and Genomes. Adjusted p-value (corrected for multiple hypothesis testing with the Benjamin–Hochberg method) < 0.05. Count: number of differentially expressed genes within a given KEGG pathway; Grey dots within plot indicate insignificant (FDR p-value cutoff = 0.05) enrichment of KEGG pathway

Host-specific microbiota members might play profound roles in maintaining basic biological processes. We next performed GO enrichment analysis of DEGs (designated as Up or Down biological processes) consisting of three parts, namely, biological process, cellular component, and molecular function (**Figure S4-6**). We selected the top 10 enriched categories in the bubble plot sorted by the DEG hit ratio from the background mouse genome. A significant dysregulation was observed in HM mice in all categories analyzed compared to MM mice under NCD treatment (**Figure S4**). Specifically, amongst the perturbated biological processes and cellular components in the NCD-fed HM mouse livers, most pathways involving the highest number of genes appeared mainly involved in mitochondrial dysfunction (**Figure S4A and B**). Lastly, significant terms that emerged in the GO molecular function analysis were also detected between HM and MM mice under NCD treatment, and the top 10 were displayed in **Supplementary Figure 4**. We also observed marked shifts in basial biological processes in NCD-fed MM mice compared to the GF littermates (**Figure S5**) and in HFD-fed HM mice compared to their GF littermates (**Figure S6**). The enriched KEGG pathways were calculated from DEGs between HM, MM vs. GF mice to characterize the contribution of host-specific microbiota to the higher-level systemic functions (**Figure 5C**). In the top 10 enriched KEGG pathways, we identified that oxidative phosphorylation, ribosome, non-alcoholic fatty liver disease, Parkinson’s disease, thermogenesis, diabetic cardiomyopathy, prion disease, Hungtington’s disease, Alzheimer’s disease, and metabolic pathways were significantly enriched in NCD-fed HM mice versus MM mice. In contrast, no down biological processes in HM mouse livers were detected (**Figure 5C**). Notably, no significantly enhanced KEGG pathways were observed from DEGs between NCD-fed HM vs. GF mice. Like NCD-fed HM mice, most of the top 10 significantly enriched KEGG pathways based on DEGs upregulated in HFD-fed HM mice were immune-related functions compared to those in HFD-fed GF mice (**Figure 5C**). KEGG pathways were also significantly enriched based on DEGs elevated in NCD-fed GF mice compared to NCD-fed MM mice, suggesting a profound transcriptional dysregulation without the presence of host-specific microbiota (**Figure 5C**).

These RNA-seq results indicate marked shifts in hepatic gene transcription, mitochondrial dysfunction, oxidative phosphorylation, intracellular signaling, and metabolism in HM mice due to exogenous microbiota colonization, revealing a profound transcriptional dysregulation. Taken together, these results indicate that a large set of profound transcripts in the liver were discriminable between mouse recipients associated with host-specific and exogenous microbiota, especially under NCD feeding conditions, and that the presence of host-specific microbial communities can profoundly maintain liver metabolic homeostasis.

### Host-microbiota mismatch feature results in microbiota dysmetabolism compared to the host-specific microbiota

The disordered production of short-chain fatty acids (SCFAs) and secondary bile acids (BAs) by the intestinal microbial communities was associated with host dysmetabolism ^16^. To evaluate if these compositional and diversity changes induced by the host-microbiota mismatch feature have microbial metabolism functional implications, we assessed the SCFAs and secondary BAs concentration in the cecal luminal samples in our mice model with host-specific or nonspecific microbiota compared to GF counterparts. The principal component analysis (PCA) showed that the carbohydrate and lipid metabolism pathway, including SCFAs and secondary BA biosynthesis, significantly differed between mice with host-specific (MM) or nonspecific microbiota (HM) (**Figure 6A**). The butyrate and acetate were significantly lower in the cecal luminal samples of HM mice than in MM mice on NCD (**Figure 6B**). Especially, host-microbiota mismatch changes the SCFA pool by decreasing butyrate to nearly undetectable levels under NCD treatment, similar to GF mice. Consistent with the metabolites on NCD treatment, butyrate, and acetate were significantly reduced in the cecal luminal samples from the HM group compared to the MM group when both were fed on HFD while displaying a similar pattern with GF mice on the same diet (**Figure 6B**). Notably, succinate was significantly increased in the cecal luminal samples from the HM group compared to the MM group fed on NCD; both were significantly higher than that in GF mice on NCD or HFD (**Figure 6B**).

**Fig. 6:**
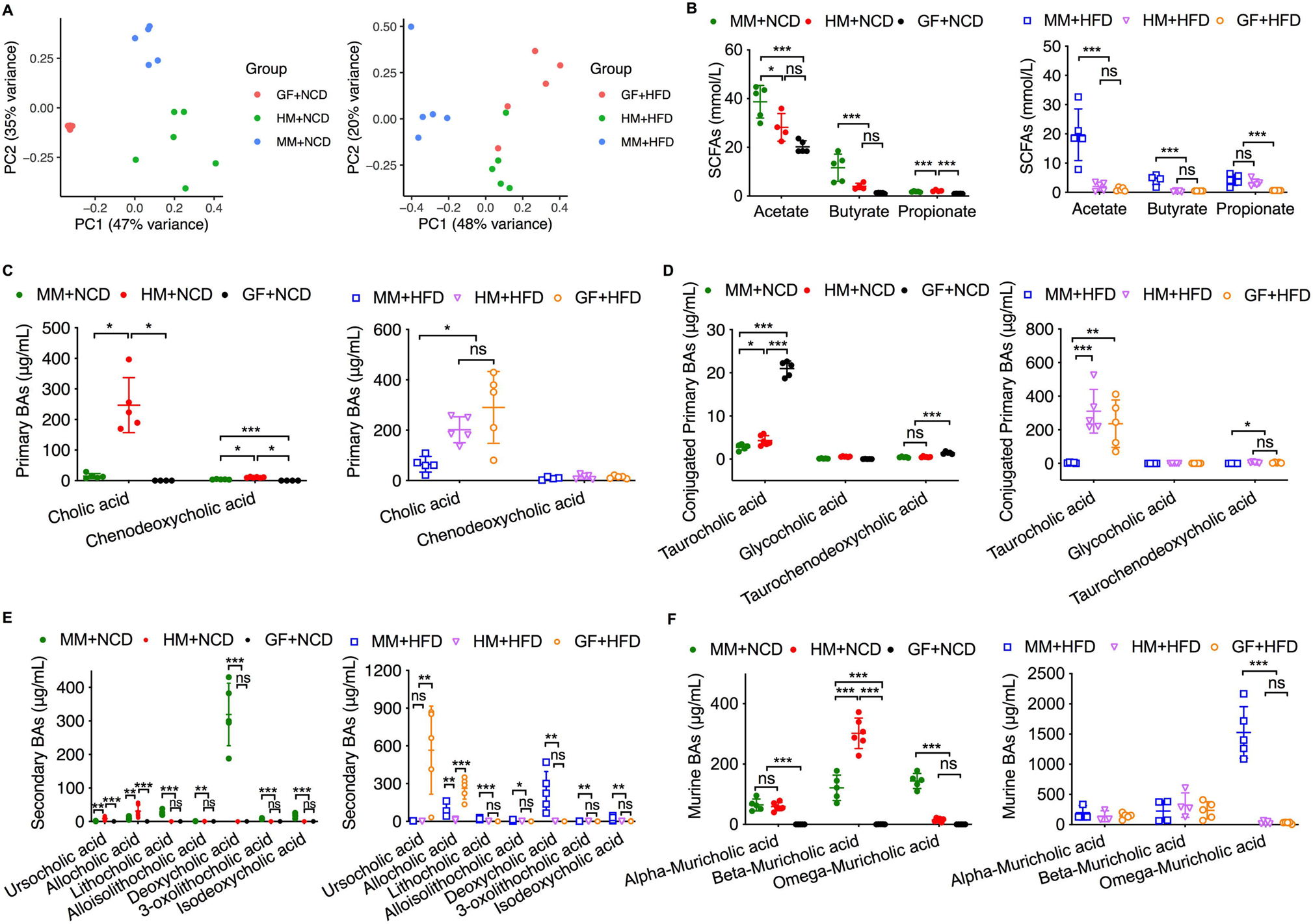
Host-specific microbiota affects SCFA and BA profiles in the cecum. **A**. PCA analysis of the cecum SCFA and BA of NCD-fed or HFD-fed HM, MM, and GF mice. **B**. Absolute quantification of the level of different SCFA in the cecal content of NCD-fed or HFD-fed HM, MM, and GF mice. **C-F**. Absolute quantification of primary (**D**), conjugated primary (**E**), and secondary (**F**) and murine BAs (**G**) in the cecal content of NCD-fed or HFD-fed HM, MM, and GF mice. Results are expressed as the mean ± SEM. n = 3–6 male mice per group. Statistical comparison was performed by testing normality using Kolmogorov–Smirnov test and then ANOVA or Kruskal–Wallis test with Turkey’s post hoc test. *p < 0.05. *p ≤ 0.05, **p ≤ 0.01, and ***p ≤ 0.001. ns, not significant.

As to the luminal bile acid profile of mice harboring host-specific or nonspecific gut microbiota, we found that host-microbiota mismatch led to remarkable changes in bile acids composition in the cecum; these alterations were characterized by significantly elevated concentrations of primary bile acid conjugates, as opposed to secondary conjugates (**Figure 6C-F**). Specifically, cecal bile acids of HM mice showed an increased proportion of many primary BAs, such as cholic acid and its derivative taurocholic acid, displaying a similar pattern with GF mice on the same diet (**Figure 6C** and **D**). Furthermore, HM tended to further dysregulate bile acid composition in the cecum with significantly lower amounts of secondary BAs (lithocholic acid, deoxycholic acid, isodeoxycholic acid, and 3-oxolithocholic acid, etc.) in cecal luminal samples from HM recipients (**Figure 6E** and **F**) compared to MM on the same diet. These differences between HM and MM were associated with diet components. Notably, HM mice exhibited bile acid dysmetabolism similar to the GF controls on either NCD or HFD treatment (**Figure 6C-F**). Thus, the host-microbiota mismatch is accompanied by significant changes in luminal SCFAs and BAs, which might affect host signaling through modulating luminal microbial metabolites, leading to whole-body metabolic homeostasis dysfunction.

### Host-microbiota mismatch feature results in gut metabolite signaling dysfunction compared to the host-specific microbiota

To evaluate if these gut secondary metabolite changes induced by the host-microbiota mismatch feature have functional implications, we assessed the host signaling in the gut samples in our mice model with host-specific or nonspecific microbiota compared to GF counterparts. HM affected gut metabolite signaling by activating both SCFA and bile acid receptors. For example, GPR109 and GPR41 transcripts were significantly higher in MM mice under NCD or HFD diet conditions compared to HM or GF counterparts (**Figure 7A**). However, these SCFA receptors were also significantly higher in HM mice compared to GF control mice under NCD diet condition, exhibiting no different levels as GF mice under HFD treatment (**Figure 7A**), which indicated that HM mice exhibited mild SCFA signaling under NCD condition. Furthermore, HM mice displayed significantly lower expression levels of the BA receptor, FXR, compared to MM mice under HFD treatment (**Figure 7B**). Interestingly, there was no significant difference in the FXR expression levels among mice on NCD (**Figure 7B**). These cecum gene expression changes suggest that HM induces gut microbiota metabolism insufficiency, alters metabolites signaling, and thus affects hepatic and systemic metabolism, due to the defect of microbiota in a mismatched host gut.

**Fig. 7:**
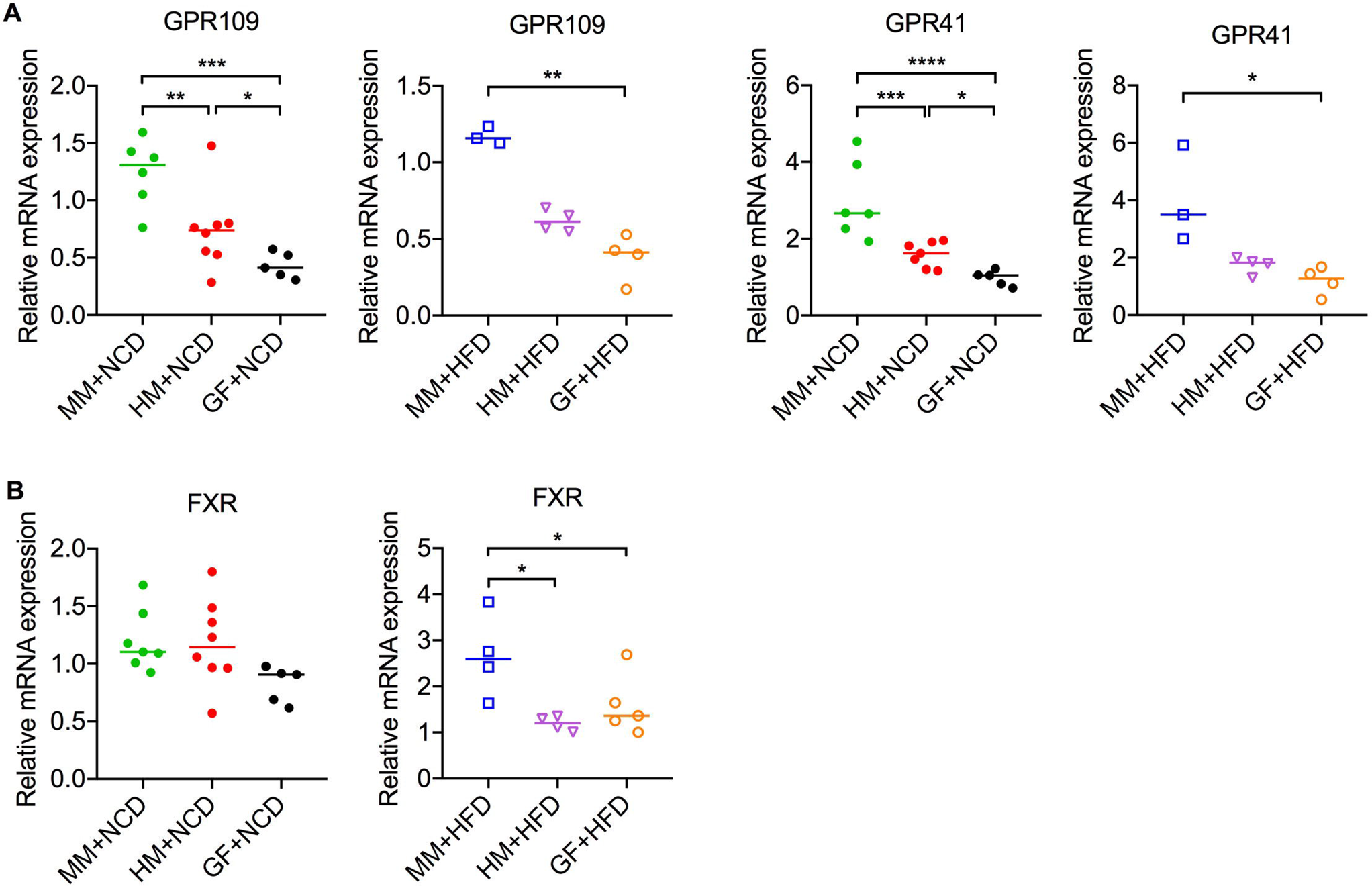
Host-specific microbiota affects SCFA and BA receptors in the cecum. **A**. Expression of the GPR109 and GPR41 genes in the cecum after 8 weeks of NCD or HFD. **B**. Expression of the FXR gene in the cecum after 8 weeks of NCD or HFD. Results are expressed as the mean ± SEM. n = 3–6 male mice per group. Statistical comparison was performed by testing normality using Kolmogorov–Smirnov test and then ANOVA or Kruskal–Wallis test with Turkey’s post hoc test. *p < 0.05. *p ≤ 0.05, **p ≤ 0.01, and ***p ≤ 0.001. ns, not significant.

## Discussion

The metabolic homeostasis of the mammal hosts is contributed by the balance among diet, host genetics, and gut microbiota. In this study, we found that a host-specific microbiota was required for maintaining the host metabolic homeostasis, while host-microbiota mismatch resulted in metabolic insufficiency. Interestingly, this metabolic insufficiency was due to the low colonization rate of exogenous microbes in the mouse gut and could be further induced by the metabolite dysfunction via those few exogenous microbes that did successfully colonize. This study provides an integrated view of the most relevant research questions concerning the fitness benefits of the mammal microbiome on the host physiology and building a better probiotic in the microbiome-targeted therapeutic.

In this study, we explored how the host responded to diet and a foreign microbiome using GF mice models associated with host-specific microbiota (mouse microbiota (MM)) or exogenous microbiota (human microbiota (HM)), with GF mice as control. We observed that MM and HM animals have very different effects on host metabolic profiles according to the observed metabolic phenomenon and metabolic rate. The exogenous microbial community (HM) did not show as detrimental effects as the indigenous one (MM) on the body weight even under HFD treatment in male mice. HM partly masked HFD’s adverse metabolic effects on the host in male mice, exhibiting a similar phenotype trend to GF mice (i.e., hypermetabolism, metabolic insufficiency, and inflexibility). The reason perhaps is because MM was more efficient at harvesting nutrients from the diet and therefore gained more weight than the mismatched group. The host-specific microbiome, in turn, promoted weight gain and metabolic dysregulation in the host under the high-fat diet and created a feed-forward loop that promotes obesity. Furthermore, metabolic cage analysis of mice associated with a different microbiome status at the end of the trial on both normal chow and high-fat diet conditions revealed that the presence of host-specific microbiota had a dominant role in shaping the host global metabolic profile. In a previously GF mouse with a host-specific microbiome on high-fat diet, consistent with the development of murine equivalent of obesity, we observed metabolic rate towards a profile resembling that found in extreme obesity. However, female mice responded to the diet more slowly than the male mice and had a different pattern of metabolic rate, consistent with hormonal impacts on the metabolic function ^17^. Long-term adaptation to both diets in females compared to males is required in future studies. We can speculate that the host-specific microbiota allows stricter control of metabolic homeostasis (e.g., metabolic rate), which, along with the reestablishment of gut ecology supports the recovery of the initial microbiota composition ^1,2^.

This metabolic insufficiency in the microbial mismatch might be due to the low colonization rate of exogenous microbes in the mouse gut. Kinetic analysis of the evolution of the microbiome over time on different hosts revealed that gut ecology and evolution have a dominant role in shaping the host bacterial profile ^1,2^. We observed that the host-microbiota mismatch resulted in remarkedly defects of microbiota, including reduced diversity and decreased abundance of specific bacterial groups (especially Firmicutes and Bacteroidetes species), which lost beneficial actions on the host. The decreased colonization rate of human microbiota in naïve mouse gut is consistent with previous studies ^18,19^. A reduction in “beneficial” flora is associated with the onset of hypermetabolism, metabolic insufficiency, and inflexibility, a hallmark predisposing factor for metabolic dysfunction. Along these lines, germ-free animals reconstituted with a foreign microbiota from human donors exacerbated metabolic alterations in the system and locally with connections to metabolic dysfunction. The mammal gut microbiome likely consists of a mix of beneficial, commensal, and pathogenic microbes ^20^. Host-specific microbiome may promise individuals to make effective use of the food source and pave the way for genomic adaptation that would promote microbiota interactions (co-metabolism and cross-feeding loop) and improve the utilization of the food resource ^21^. The congruent relationships between hosts and these unique and specific host-specific community members are likely maintained through the microbiome’s efficiently positive effects on host performance and fitness.

To further determine the mechanistic basis by which host-microbe mismatch impacts host metabolism locally, we performed liver transcriptomic analysis on the host and metabolites analysis on the microbiota. Our findings indicated that the host-specific microbiota might play critical roles in coordinating the interaction between diet and host genetics in the liver. Gene expression and metabolism were altered in GF and humanized mice. Indeed, among the gene pathways in the liver that were significantly modulated by the host-microbe mismatch were those involved in mitochondrial dysfunction, oxidative phosphorylation, intracellular signaling, and metabolism. Specifically, most pathways involving the highest number of genes appeared mainly involved in mitochondrial dysfunction in the NCD-fed HM mouse livers. Mitochondria, the powerhouses of our cells, play an essential role in the process of health, disease, and aging and mitochondrial dysfunction has been implicated in nearly all pathologic and toxicologic conditions ^22,23^. Increased mitochondrial activity or mitochondrial dysfunction is widely accepted as one of the metabolic characteristics in GF and humanized mice ^24,25^. Consistent with the dysregulated immune profiles observed in these diseases, the enriched KEGG pathways also confirmed a profound immune-related transcriptional dysregulation, e.g., increased oxidative phosphorylation, ribosome, non-alcoholic fatty liver disease, Parkinson’s disease, thermogenesis, diabetic cardiomyopathy, prion disease, Hungtington’s disease, and Alzheimer’s disease pathways without the presence of host-specific microbiota. The presence of host-specific microbial communities can profoundly maintain liver metabolic homeostasis, while an experimentally mediated disruption of phylosymbiosis will have functional costs that may lower host fitness or performance in the liver. Overall, host-microbiome mismatch resulted in harmful consequences in the liver transcription because of metabolic dysfunction. Strikingly, high-fat diet had a greater impact on the liver transcriptome than either microbiota. We hypothesized that activation of cell signaling pathways by microbiota-derived molecules during the host-specific microbiota colonization is crucial for host physiological maturation and downstream metabolism. In future studies, the signals from that provided by the microbiome responsible for this phenomenon need to be characterized.

Metabolite molecular mechanisms provided by the mismatched microbiome may contribute to the host resulting in metabolic dysfunction ^12,13^. Recently, the gut microbial metabolites were suggested to regulate host fatty acid and lipid metabolic homeostasis, through SCFA and bile acid signaling ^11,15^. In our study, the exogenous microbes could not meet the host metabolic needs for butyrate as the main fuel source of the colon and thus, which further affected whole-body metabolic homeostasis by lowering the SCFA signaling GPR109 and GPR41 in the cecum. Our study adds to this body of knowledge by demonstrating that host-microbiota mismatch decreased levels of SCFAs, as the colon’s primary source of energy, likely promotes energy demand by the colon ^14^. In addition to SCFAs, the host-microbiota mismatch also caused bile acid dysmetabolism and its downstream targets (e.g., FXR) in the cecum. Bile acids circulate in the intestine where primary bile acids are further deconjugated and converted into secondary bile acids by the microbiota and become further mediators in glucose metabolism ^15^. In this study, we observed that host-microbiota mismatch significantly increased the concentration of taurine-conjugated bile acids ^7,26^ and decreased the proportion of secondary bile acids in the cecum. Additionally, unconjugated bile acids, such as cholic acid and chenodeoxycholic acid, are strong agonist for bile acid receptors, including FXR signaling, these receptors activates transcriptional networks and signaling cascades relevant for cholesterol and lipid metabolism, maintenance of glucose, and hepatic homeostasis ^17^. Our findings showed that FXR expression was downregulated in the cecum in host-microbiota mismatched mice under the HFD condition, all hallmarks described in GF or antibiotic-treated mice mice ^12,13^. Furthermore, FXR deactivation in the gut suggests an increased uptake of BAs and consequently, increases the de novo synthesis of primary BAs in hepatocytes. Moreover, the pro-inflammatory properties of primary bile acids have been described ^15,27^. Hence, host-microbiome mismatch induces changes in BA physiology and could likely affect host metabolism through various methods, most of which remain to be explored. Lastly, we observed some of the hallmarks of dysmetabolism of HM-associated mice at an intermediate level between the MM mice and GF mice, which suggested that these colonized exogenous microbes can still perform metabolic function though at a very limited level. Altogether, these data suggest that host-microbe mismatch’s impact on SCFA and bile acid metabolism may be one of the mechanisms by which the exogenous microbiome potentiates HFD-induced hepatic and whole-body metabolic impairment and host dysfunctions. These results defined a novel mechanism by which the host-specific microbiome regulates host metabolic homeostasis by microbial metabolites.

The host-associated microbiome can generate and contribute to the host plastic responses responding to the diet shift ^28^. In our study, we demonstrated that within the host-specific microbial community, the diets could characterize the high variation of microbiome composition and abundance; these microbial dynamics could potentially influence many dimensions of the host’s phenotype from that provided by microbial fitness. These host-specific microbiome-mediated changes can be induced by the environment (i.e., diet) and thus produce effects equivalent to what is traditionally considered host phenotypic plasticity. In contrast, both the host and microbiota plastic responses responding to the diet shift for an exogenous microbiome (i.e., the human microbiome) is quite limited. It seems like that the molecular signal communication (both positive and negative signals) between host-microbe is better when matched through a similar language, while, GF mice aren’t transducing any environmental signals whatsoever and the HM aren’t transducing as sufficient or same signals.

We demonstrated that host-microbiota mismatch, like microbial dysbiosis, augments some of the deleterious host metabolic effects by modifying multiple targets (i.e., altering the diversity and functional potential of intestinal microbes, increasing metabolic rate) and promoting dysfunction, development of heightened immune response in livers and disproportionate SCFA and bile acid concentrations in the gut lumina. All these pathways ultimately converge to further disturb the host metabolic function, especially in the HFD setting. In summary, host-microbe mismatch has significant metabolic consequences. Though it is unclear whether these metabolic consequences of host-microbe mismatch are due to the colonization selection of the microbiome or the insufficiency of metabolomics, or a combination thereof, it is very likely that they will have a significant impact on host physiological processes. Collectively, our study provides a conceptual framework to further test this hypothesis in other mammal hosts and warrants the evaluation of preventative strategies, such as probiotics use, when suppressing microbial dysbiosis-related disease.

## Conclusion

Our study suggests that the host-specific microbiota is critical for maintaining the metabolic homeostasis of the host, and gut microbiota manipulating in disease treatment are more host-specific than previously understood. The important signatures of the host-specific microbiota we discovered are the home-site advantage for colonization and the sufficient and proper metabolites it supplied. Altogether, our multi-omics study is the first report on host-specific microbiota in metabolic homeostasis. Given the mutual function of the host-specific microbiota in host fitness is complex and multifaceted, we believe that this study has laid a solid foundation for further investigation of the precise mechanisms involved in gut microbiota and the development of microbiota therapeutics for disease. An important future research avenue is to disentangle the various individual contributions of gut symbionts to reconstruct and understand their combined contribution to host fitness, in managed as well as a consistent set of controlled experimental and bioinformatic approaches. Another challenging question is the extent to which the environment contributes to gut microbiome composition and dynamics/evolution, and how this, in turn, affects host fitness. Although mouse models are popular in host microbiota-metabolic disease studies due to their relatively low cost and easy background control, studies using mouse models should interpret their results cautiously. Translational microbiome research and therapeutic development must account for host-specific engraftment in fecal microbiota transplantation and probiotics. We argue, to achieve these goals, standardized research methods and analyses based on the ecological and evolutionary perspective are required, as well as communal resources that integrate information from disparate studies.

## Methods

### Animal husbandry

All animal studies have been approved by the institutional animal care and use committee at the University of Chicago. All mice used for these studies were on a C57Bl/6J genetic background. Germ-free (GF) mice (the GF animal facility of the University of Chicago) were bred under the GF rodent breeding conditions at the University of Chicago. The GF status of the mice was assessed every week using aerobic and anaerobic cultivation and 16S PCR of mouse feces to confirm the absence of bacteria, molds, and yeast throughout the experiment. Specific Pathogen Free (SPF) mice were bred within the University of Chicago SPF barrier facility. Unless indicated otherwise, animals were fed an autoclaved standard diet, low in fat and rich in plant polysaccharides ad libitum. Mice received microbiota transplants at 3∼4 weeks of age. The bedding was replaced in all experiments every 7 days. All the mice were maintained on a 12:12-h lighted-dark cycle. The mice were treated with HM or MM content by single oral gavage. Mice were euthanized, and tissue samples were harvested at the trial’s end. Briefly, the cecal contents were freshly frozen and stored at −80°C for subsequent assays. The high-fat diet (18% human milk fat) used in the study was customized and purchased from Envigo (TD.97222).

### Metabolic phenotyping

For energy metabolism analysis, mice were housed individually and monitored in germ-free Promethion metabolic cage systems (Sable Systems International, Las Vegas, USA) following diet treatment. Metabolic cages are located within an ambient temperature and light-controlled environmental enclosure. Mice were exposed to a 12 h light/ dark cycle and ambient temperature of 22°C. They were housed on pine chip bedding without nestlets and provided ad libitum access to NCD or HFD and water. Mice were housed in the metabolic cages for 1 week under the original diet treatment, where gas exchange, energy expenditure, and total activity (all meters/pedometers) were continuously measured. Oxygen consumption (VO_2_), carbon dioxide production (VCO_2_), respiratory exchange ratio (RQ), and energy expenditure (EE), were measured every 5 min intervals for 30 seconds for analysis (12 h dark-light phase comparison). The respiratory exchange quotient (RQ) was calculated as a ratio of CO_2_ production over O_2_ consumption and was used as an indicator of substrate utilization. An RQ of >0.8 indicates primarily carbohydrate metabolism, 0.8 mixed carbohydrate and fat metabolism, and <0.8 primarily fat metabolisms. Energy expenditure was calculated using the Weir equation ^29^: Kcal/h = 60 × (0.003941 × VO_2_ + 0.001106 × VCO_2_), where VO_2_ is O_2_ consumption and VCO_2_ is CO_2_ output. Food and water intake, along with body mass, are assessed gravimetrically. Ambulatory activity is determined simultaneously every second with the collection of the calorimetry data. Ambulatory activity and position are detected with XYZ beam arrays (BXYZ-R; Sable Systems, Las Vegas, NV) with a beam spacing of 1.0 cm interpolated to a centroid resolution of 0.25 cm. MetaScreen v2.2.18 coordinated data acquisition and instrument control, and the raw data were processed using ExpeData v1.7.30 (Sable Systems).

### Liver histology

The liver sample from each mouse was formalin-fixed, embedded in paraffin, and sectioned (5 μm), followed by deparaffination for hematoxylin-eosin (H&E) stain using standardized protocols, and scored for severity of steatosis and inflammation by an experienced pathologist blinded to the experiments. A semi-quantitative system of scoring the features of nonalcoholic fatty liver disease (NAFLD) called the NAFLD Activity Score (NAS) was developed based on the histological assessment by routine stains. Total NAS represents the sum of scores for steatosis (0–3), lobular inflammation (0–3), ballooning (0–2), and ranges from 0-8. Steatosis scores: percentage of surface area involved by steatosis as evaluated on low to medium power examination; 0 (0–5%), 1 (5–33%), 2 (33–66%), 3 (>66%). Lobular Inflammation scores: 0 (None), 1 (< 2 focis / 200x), 2 (2-4 focis / 200x), 3 (> 4 focis / 200x). Hepatocyte Ballooning scores: 0 (None), 1 (Few balloon cells), 2 (Many cells/prominent ballooning) ^30^.

### Epididymal adipose tissue histology

Adipose tissue morphometric analysis was performed on the sections stained by hematoxylin-eosin-saffron (HES stain). Slides were digitalized using a Panoramic digital slide scanner (3DHistech Ltd., Budapest, Hungary), and adipocyte morphometry was analyzed, quantified, and photographed using Panoramic Viewer software by 3DHistech. The number of adipocytes per microscopic field (at least 5 fields per tissue) was determined at a magnification of 200x, and the average surface area of the adipocytes (in square micrometers) was calculated.

### DNA isolation and 16S ribosomal RNA (rRNA) gene sequencing

DNA was extracted using the QIAamp PowerFecal Pro DNA kit (Qiagen). Before extraction, samples were subjected to mechanical disruption using a bead beating method. Briefly, samples were suspended in a bead tube (Qiagen) along with lysis buffer and loaded on a bead mill homogenizer (Fisher brand). Samples were then centrifuged, and the supernatant was resuspended in a reagent that effectively removed inhibitors. DNA was then purified routinely using a spin column filter membrane and quantified using Qubit. V4-V5 region within the 16S rRNA gene was amplified using universal bacterial primers – 563F (5’-nnnnnnnn-NNNNNNNNNNNN-AYTGGGYDTAAA-GNG-3’) and 926R (5’-nnnnnnnn-NNNNNNNNNNNN-CCGTCAATTYHT-TTRAGT-3’), where ‘N’ represents the barcodes, ‘n’ are additional nucleotides added to offset primer sequencing. Approximately ∼412bp region amplicons were then purified using a spin column-based method (Minelute, Qiagen), quantified, and pooled at equimolar concentrations. Illumina sequencing-compatible Unique Dual Index (UDI) adapters were ligated onto the pools using the QIAseq 1-step amplicon library kit (Qiagen). Library QC was performed using Qubit and Tapestation and sequenced on the Illumina MiSeq platform to generate 2×250bp reads.

### 16S rRNA gene pyrosequencing data preprocessing and analysis

We used DADA2 algorithm (v1.18.0) ^31^, as our default pipeline for processing MiSeq 16S rRNA reads with minor modifications in R (v4.0.3). Specifically, reads were first trimmed at 180 bp for both forward and reverse reads to remove low-quality nucleotides. Chimeras were detected and removed using the default consensus method in the dada2 pipeline. Then, exact sequence variants (ASVs) with lengths between 300 bp and 360 bp were kept and deemed as high-quality ASVs. Taxonomy of the resultant ASVs was assigned to the genus level using the RDP database with a minimum bootstrap confidence score of 50. Multiple sequence alignment of all ASVs was performed to generate a neighbor-join phylogenetic tree using the R package “MSA” (v1.22.0) and “ape” (v5.6.1). Alpha and beta-diversity analyses were performed in R using the *phyloseq* package ^32^. Alpha diversity was calculated by Shannon’s diversity index ^33^. Principal coordinate analysis (PCoA) was performed based on weighted and unweighted UniFrac distances, a method for computing differences between microbial communities based on phylogenetic information ^34^. Weighted UniFranc considered both ASVs presence and absence and abundance distances, and unweighted UniFrac only considered ASVs presence. Permutational multivariate analysis of variance (PERMANOVA, R function adonis (vegan, 999 permutations)) was used to analyze paired statistical differences in beta diversity ^35^. Benjamini–Hochberg false discovery rate (FDR) correction was used to correct for multiple hypothesis testing ^36^. Significantly different ASVs were determined using the analysis platform of STAMP.

### Liver RNA sequencing and data preprocessing, processing, normalization, and differential expression analysis

The mouse liver samples were collected from MM and HM-associated mice after 8 weeks of dietary exposure. Five to six replicates were taken for each distinct condition. Frozen liver tissue was processed to isolate total RNA as previously described ^37^. The RNA samples were sequenced on an Illumina NovaSEQ6000 platform via the genomics facility at the University of Chicago. The samples were sequenced by RNA-seq technology to produce raw reads in fastq format. The raw fastq files generated from the RNA-seq analysis were examined by the fastqc to evaluate the quality. A batch correction was done using ComBat-Seq ^38^ from R library SVA to remove the unwanted technical variation from the dataset from the two sequencing runs. The STAR ^39^ (version 2.4.2a, Stanford University, CA) aligner was used to map the raw reads to the reference mouse genome (GRCm38 https://www.ncbi.nlm.nih.gov/grc). The STAR default parameter for the maximum mismatches is 10, which is optimized based on mammalian genomes and recent RNA-seq data. The genetic features from Gencode ^40^ vM23 were extracted from the resulting bam file produced by STAR. The raw gene expression count matrix was then generated by featureCounts ^41^ (version subread-1.4.6-p1). We used the EdgeR ^42^ package (version 3.22.5) to filter low-expressed genes where genes expressed below ten counts-per-million (CPM) in 4 samples were removed. The Limma-Voom ^43^ pipeline (version 3.38.3) was applied to normalize the filtered count matrix and perform differential gene expression analysis for pairwise comparisons. We further focused on the impact on gene expression profiles by diet changes and longitudinal variations. Differentially expressed genes (DEGs) were identified using FDR adjusted P-value cutoff of 0.05. Enrichment analysis on significant DEGs with mouse-specific GO terms and KEGG pathways was conducted through Lynx enrichment ^44^. Significantly enriched categories were selected using FDR adjusted P-value < 0.05.

Multidimensional scaling (MDS) plot was used to show variation among samples based on the normalized RNA-seq data. Each point represents 1 sample, and the distance between 2 points reflects the corresponding samples’ leading logFC (Folder change). The leading logFC was the average of the largest absolute logFC between each pair of samples. We labeled and colored the samples by the microbiome and diet group. We applied a volcano plot of DEGs in various contrast groups. The logFC for each gene was plotted against the FDR-adjusted P-value. The horizontal red line represented a significance threshold of 0.05. The colored points distinguished the negative or positive logFC within the contrast groups. The Venn diagrams were used to show the overlapping DEGs upregulated in two related contrast groups. The heatmap was generated by ComplexHeatmap ^45,46^, venn diagram was by VennDiagram ^47^, and MDS was from edgeR ^42^. We showed the top 10 enriched categories and the top 10 KEGG pathways in the bubble plot sorted by the DEG hit ratio from the background mouse genome. We considered all the three sub-ontology categories (BP for Biological Process, MF for Molecular Function, and CC for Cellular Component) for the enriched categories.

### Metabolomics

The cecal metabolome was analyzed across three mass spectrometry platforms to capture quantitative and qualitative levels of gut-derived metabolites with varying physicochemical properties, such as hydrophobicity, size, and charge, by the DFI Host-Microbe Metabolomics Facility (DFI-HMMF) at the University of Chicago with the proposed methods and analysis pipelines.

#### SCFAs and bile acids Extraction from Cecal Material

Extraction solvent (80% methanol spiked with internal standards and stored at −80 °C) was added to pre-weighed fecal/cecal samples at a ratio of 100 mg of material/mL of extraction solvent in beadruptor tubes (Fisherbrand; 15-340-154). Samples were homogenized at 4 °C on a Bead Mill 24 Homogenizer (Fisher; 15-340-163), set at 1.6 m/s with 6 thirty-second cycles, 5 seconds off per cycle. Samples were then centrifuged at −10 °C, 20,000 x g for 15 min and the supernatant was used for subsequent metabolomic analysis.

### SCFA Analysis using GC-nCI-MS and Derivatization

Short chain fatty acids were derivatized as described by Haak et al. with the following modifications.1 The metabolite extract (100 µL) was added to 100 µL of 100 mM borate buffer (pH 10) (Thermo Fisher, 28341), 400 µL of 100 mM pentafluorobenzyl bromide (Millipore Sigma; 90257) in Acetonitrile (Fisher; A955-4), and 400 µL of n-hexane (Acros Organics; 160780010) in a capped mass spec autosampler vial (Microliter; 09-1200). Samples were heated in a thermomixer C (Eppendorf) to 65 °C for 1 hour while shaking at 1300 rpm. After cooling to RT, samples were centrifuged at 4 °C, 2000 x g for 5 min, allowing phase separation. The hexanes phase (100 µL) (top layer) was transferred to an autosampler vial containing a glass insert and the vial was sealed. Another 100 µL of the hexanes phase was diluted with 900 µL of n-hexane in an autosampler vial. Concentrated and dilute samples were analyzed using a GC-MS (Agilent 7890A GC system, Agilent 5975C MS detector) operating in negative chemical ionization mode, using a HP-5MSUI column (30 m x 0.25 mm, 0.25 µm; Agilent Technologies 19091S-433UI), methane as the reagent gas (99.999% pure) and 1 µL split injection (1:10 split ratio). Oven ramp parameters: 1 min hold at 60 °C, 25 °C per min up to 300 o C with a 2.5 min hold at 300 °C. Inlet temperature was 280 °C and transfer line was 310 o C. A 10-point calibration curve was prepared with acetate (100 mM), propionate (25 mM), butyrate (12.5 mM), and succinate (50 mM), with 9 subsequent 2x serial dilutions. Data analysis was performed using MassHunter Quantitative Analysis software (version B.10, Agilent Technologies) and confirmed by comparison to authentic standards. Normalized peak areas were calculated by dividing raw peak areas of targeted analytes by averaged raw peak areas of internal standards.

#### Bile Acid Analysis

Bile acids were analyzed using LCMS. The metabolite extract (75 μL) was added to prelabeled mass spectrometry autosampler vials (Microliter; 09-1200) and dried down completely under a nitrogen stream at 30 L/min (top) 1 L/min (bottom) at 30 °C (Biotage SPE Dry 96 Dual; 3579M). Samples were resuspended in 50:50 Water: Methanol (750 μL). Vials were added to a thermomixer C (Eppendorf) to resuspend analytes at 4 °C, 1000 rpm for 15 min with an infinite hold at 4 °C. Samples were then transferred to prelabeled microcentrifuge tubes and centrifuged at 4 °C, 20,000 x *g* for 15 min to remove insoluble debris. The supernatant (700 μL) was transferred to a fresh, prelabeled mass spectrometry autosampler vial. Samples were analyzed on a liquid chromatography system (Agilent 1290 infinity II) coupled to a quadrupole time-of-flight (QTOF) mass spectrometer (Agilent 6546), operating in negative mode, equipped with an Agilent Jet Stream Electrospray Ionization source. The sample (5 μL) was injected onto an XBridge© BEH C18 Column (3.5 μm, 2.1 x 100 mm; Waters Corporation, PN) fitted with an XBridge© BEH C18 guard (Waters Corporation, PN) at 45 °C. Elution started with 72% A (Water, 0.1% formic acid) and 28% B (Acetone, 0.1% formic acid) with a flow rate of 0.4 mL/min for 1 min and linearly increased to 33% B over 5 min, then linearly increased to 65% B over 14 min. Then the flow rate was increased to 0.6 mL/min and B was increased to 98% over 0.5 min and these conditions were held constant for 3.5 min. Finally, re-equilibration at a flow rate of 0.4 mL/min of 28% B was performed for 3 min. The electrospray ionization conditions were set with the capillary voltage at 3.5 kV, nozzle voltage at 2 kV, and detection window set to 100-1700 *m/z* with continuous infusion of a reference mass (Agilent ESI TOF Biopolymer Analysis Reference Mix) for mass calibration. A ten-point calibration curve was used for quantitation. Data analysis was performed using MassHunter Profinder Analysis software (version B.10, Agilent Technologies) and confirmed by comparison with authentic standards. Normalized peak areas were calculated by dividing raw peak areas of targeted analytes by averaged raw peak areas of internal standards.

### RNA isolation and quantitative RT-PCR

Frozen liver and cecum scraping tissues were processed to isolate total RNA using a Trizol extraction method as previously described ^37^. Reverse transcription to cDNA was performed using SuperScript II (invitrogen) and random hexamers via the Roche Transcriptor High Fidelity cDNA Synthesis Kit (1ug total RNA for each sample). Forward and Reverse primers were added to iQSYBR Green PCR Supermix (Bio-rad) to amplify *gpr109* (Fwd: GGCGTGGTGCAGTGAGCAGT; Rev: GGCCCACGGACAGGCTAGGT), *gpr41* (Fwd: TTCTGAGCGTGGCCTATCCA; Rev: AGACTACACTGACCAGACCAG), and *fxr* (Fwd: CTTGATGTGCTACAAAAGCTGTG; Rev: ACTCTCCAAGACATCAGCATCTC). Real-time quantitative PCR was performed on a Bio-Rad CFX Maestro System. Real-time quantitative PCR was performed using the Biorad CFX Maestro qPCR system. Gene expression was normalized to the housekeeping gene *gapdh*. Data were presented as fold changes as compared to MM mice under each diet condition.

### Statistical Analysis

Statistical analysis was performed using the PRISM statistical software package ^48^. Data are presented as means +/− standard error of the mean (SEM), with statistical significance set at p< 0.05, 0.01, or 0.001. Statistical significance was determined via the nonparametric Kruskal-Wallis test followed by Turkey’s multiple comparisons, where p<0.05 was considered significant.

## Acknowledgments

The authors are grateful to the Animal Resources Center of the University of Chicago for managing the SPF and GF mice lines (The University of Chicago, Chicago, IL 60637, USA). This work was funded by grants from the NIH (RC2DK122394).

## Contributions

N.F. and E.C. directed and designed the study; N.F., M.S., B.T. T.L., A.T, and S.M., managed and performed the mouse work; N.F. and T.L. performed the sample analysis and interpretation. A.M.S. performed the metabolomic analysis. M. G. performed the RNA isolation and quantitative RT-PCR; B. X., N.F., and D.S. analyzed and interpreted the RNA sequencing data; N.F. analyzed the metagenomic sequencing data; and N.F., wrote the paper, All authors reviewed the manuscript and approved the final manuscript.

## Reference

1 Rawls, J. F., Mahowald, M. A., Ley, R. E. & Gordon, J. I. Reciprocal gut microbiota transplants from zebrafish and mice to germ-free recipients reveal host habitat selection. Cell 127, 423–433 (2006).

2 Seedorf, H. et al. Bacteria from diverse habitats colonize and compete in the mouse gut. Cell 159, 253–266 (2014).

3 Ley, R. E. et al. Evolution of mammals and their gut microbes. science 320, 1647–1651 (2008).

4 Jovel, J., Dieleman, L. A., Kao, D., Mason, A. L. & Wine, E. The human gut microbiome in health and disease. Metagenomics, 197–213 (2018).

5 Chung, H. et al. Gut immune maturation depends on colonization with a host-specific microbiota. Cell 149, 1578–1593 (2012).

6 Tremaroli, V. & Bäckhed, F. Functional interactions between the gut microbiota and host metabolism. Nature 489, 242–249 (2012).

7 Devkota, S. et al. Dietary-fat-induced taurocholic acid promotes pathobiont expansion and colitis in Il10−/− mice. Nature 487, 104–108 (2012).

8 Bäckhed, F. et al. The gut microbiota as an environmental factor that regulates fat storage. Proceedings of the national academy of sciences 101, 15718–15723 (2004).

9 Fei, N. & Zhao, L. An opportunistic pathogen isolated from the gut of an obese human causes obesity in germfree mice. The ISME journal 7, 880–884 (2013).

10 Carvalho, B. et al. Modulation of gut microbiota by antibiotics improves insulin signalling in high-fat fed mice. Diabetologia 55, 2823–2834 (2012).

11 Canfora, E. E., Meex, R. C., Venema, K. & Blaak, E. E. Gut microbial metabolites in obesity, NAFLD and T2DM. Nature Reviews Endocrinology 15, 261–273 (2019).

12 Wichmann, A. et al. Microbial modulation of energy availability in the colon regulates intestinal transit. Cell host & microbe 14, 582–590 (2013).

13 Sayin, S. I. et al. Gut microbiota regulates bile acid metabolism by reducing the levels of tauro-beta-muricholic acid, a naturally occurring FXR antagonist. Cell metabolism 17, 225–235 (2013).

14 Tolhurst, G. et al. Short-chain fatty acids stimulate glucagon-like peptide-1 secretion via the G-protein–coupled receptor FFAR2. Diabetes 61, 364–371 (2012).

15 Ridlon, J. M., Kang, D. J., Hylemon, P. B. & Bajaj, J. S. Bile acids and the gut microbiome. Current opinion in gastroenterology 30, 332–338 (2014).

16 Agus, A., Clément, K. & Sokol, H. Gut microbiota-derived metabolites as central regulators in metabolic disorders. Gut 70, 1174–1182 (2021).

17 Mauvais-Jarvis, F. Sex differences in metabolic homeostasis, diabetes, and obesity. Biology of sex differences 6, 1–9 (2015).

18 Li, Y., Cao, W., Gao, N. L., Zhao, X.-M. & Chen, W.-H. Consistent alterations of human fecal microbes after transplantation into germ-free mice. Genomics, Proteomics & Bioinformatics 20, 382–393 (2022).

19 Li, N. et al. Spatial heterogeneity of bacterial colonization across different gut segments following inter-species microbiota transplantation. Microbiome 8, 1–24 (2020).

20 Bäumler, A. J. & Sperandio, V. Interactions between the microbiota and pathogenic bacteria in the gut. Nature 535, 85–93 (2016).

21 Chang, C.-Y., Bajić, D., Vila, J. C., Estrela, S. & Sanchez, A. Emergent coexistence in multispecies microbial communities. Science 381, 343–348 (2023).

22 Pieczenik, S. R. & Neustadt, J. Mitochondrial dysfunction and molecular pathways of disease. Experimental and molecular pathology 83, 84–92 (2007).

23 Lowell, B. B. & Shulman, G. I. Mitochondrial dysfunction and type 2 diabetes. Science 307, 384–387 (2005).

24 Bäckhed, F., Manchester, J. K., Semenkovich, C. F. & Gordon, J. I. Mechanisms underlying the resistance to diet-induced obesity in germ-free mice. Proceedings of the National Academy of Sciences 104, 979–984 (2007).

25 Hirschey, M. D. et al. SIRT3 deficiency and mitochondrial protein hyperacetylation accelerate the development of the metabolic syndrome. Molecular cell 44, 177–190 (2011).

26 David, L. A. et al. Diet rapidly and reproducibly alters the human gut microbiome. Nature 505, 559–563 (2014).

27 Fiorucci, S. et al. Bile acid signaling in inflammatory bowel diseases. Digestive Diseases and Sciences 66, 674–693 (2021).

28 Singh, R. K. et al. Influence of diet on the gut microbiome and implications for human health. Journal of translational medicine 15, 1–17 (2017).

29 Cunningham, J. Calculation of energy expenditure from indirect calorimetry: assessment of the Weir equation. Nutrition (Burbank) 6, 222–223 (1990).

30 Kleiner, D. E. et al. Design and validation of a histological scoring system for nonalcoholic fatty liver disease. Hepatology 41, 1313–1321 (2005).

31 Callahan, B. J. et al. DADA2: High-resolution sample inference from Illumina amplicon data. Nature methods 13, 581–583 (2016).

32 McMurdie, P. J. & Holmes, S. phyloseq: an R package for reproducible interactive analysis and graphics of microbiome census data. PloS one 8, e61217 (2013).

33 Peet, R. K. The measurement of species diversity. Annual review of ecology and systematics 5, 285–307 (1974).

34 Lozupone, C. & Knight, R. UniFrac: a new phylogenetic method for comparing microbial communities. Applied and environmental microbiology 71, 8228–8235 (2005).

35 Anderson, M. J. A new method for nonLJparametric multivariate analysis of variance. Austral ecology 26, 32–46 (2001).

36 Storey, J. D. A direct approach to false discovery rates. Journal of the Royal Statistical Society Series B: Statistical Methodology 64, 479–498 (2002).

37 Leone, V. et al. Effects of diurnal variation of gut microbes and high-fat feeding on host circadian clock function and metabolism. Cell host & microbe 17, 681–689 (2015).

38 Zhang, Y., Parmigiani, G. & Johnson, W. E. ComBat-seq: batch effect adjustment for RNA-seq count data. NAR genomics and bioinformatics 2, lqaa078 (2020).

39 Dobin, A. et al. STAR: ultrafast universal RNA-seq aligner. Bioinformatics 29, 15–21 (2013).

40 Frankish, A. et al. GENCODE reference annotation for the human and mouse genomes. Nucleic acids research 47, D766–D773 (2019).

41 Chen, Y., Lun, A. T. & Smyth, G. K. From reads to genes to pathways: differential expression analysis of RNA-Seq experiments using Rsubread and the edgeR quasi-likelihood pipeline. F1000Research 5 (2016).

42 Robinson, M. D., McCarthy, D. J. & Smyth, G. K. edgeR: a Bioconductor package for differential expression analysis of digital gene expression data. bioinformatics 26, 139–140 (2010).

43 Ritchie, M. E. et al. limma powers differential expression analyses for RNA-sequencing and microarray studies. Nucleic acids research 43, e47–e47 (2015).

44 Sulakhe, D. et al. Lynx: a database and knowledge extraction engine for integrative medicine. Nucleic acids research 42, D1007–D1012 (2014).

45 Gu, Z., Eils, R. & Schlesner, M. Complex heatmaps reveal patterns and correlations in multidimensional genomic data. Bioinformatics 32, 2847–2849 (2016).

46 Villanueva, R. A. M. & Chen, Z. J. (Taylor & Francis, 2019).

47 Chen, H. & Boutros, P. C. VennDiagram: a package for the generation of highly-customizable Venn and Euler diagrams in R. BMC bioinformatics 12, 1–7 (2011).

48 Swift, M. L. GraphPad prism, data analysis, and scientific graphing. Journal of chemical information and computer sciences 37, 411–412 (1997).

