## Supplementary figures and images for "The Host-specific Microbiota is Required for Diet-Specific Metabolic Homeostasis"

### Supplementary Figure 1

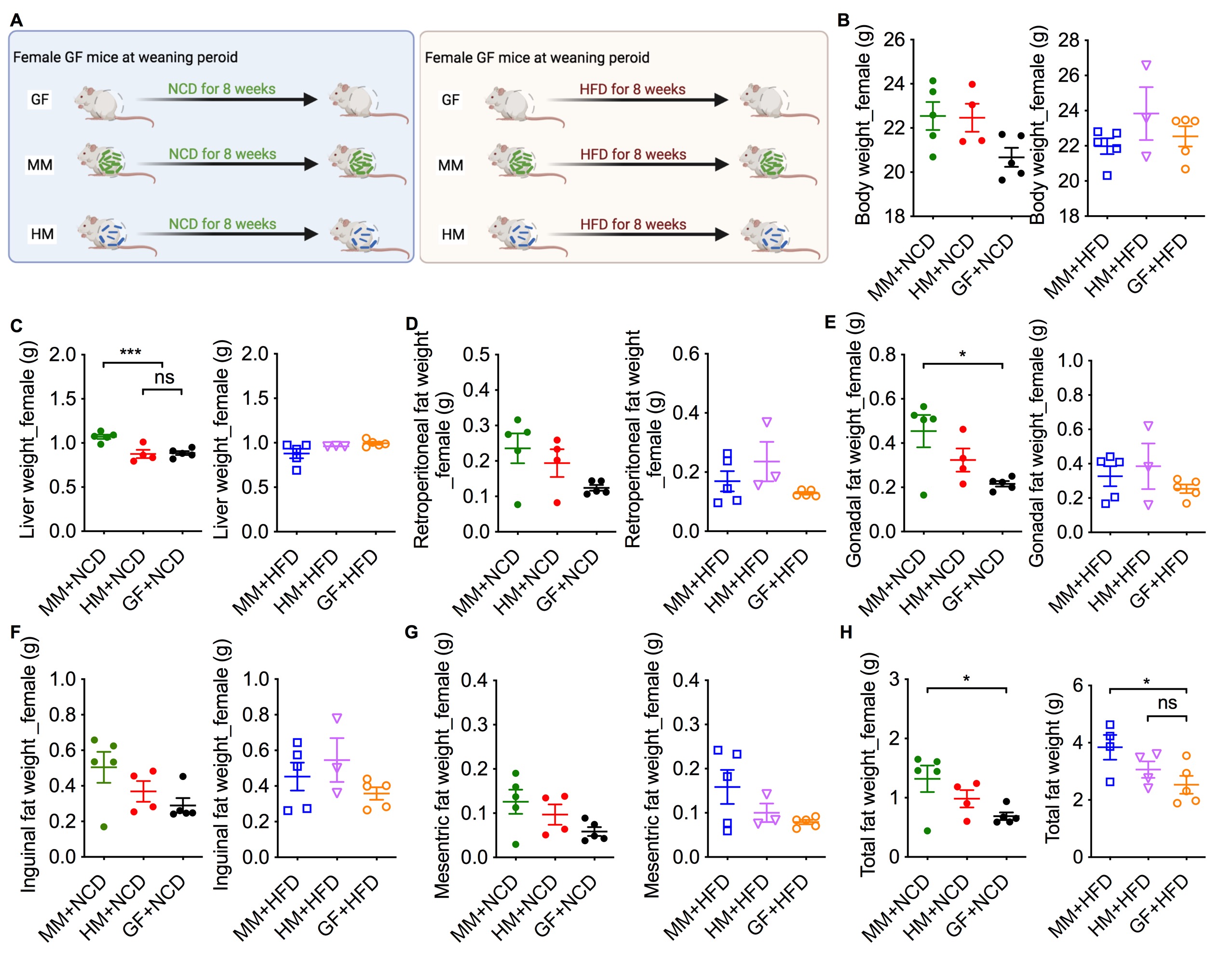

### Supplementary Figure 2

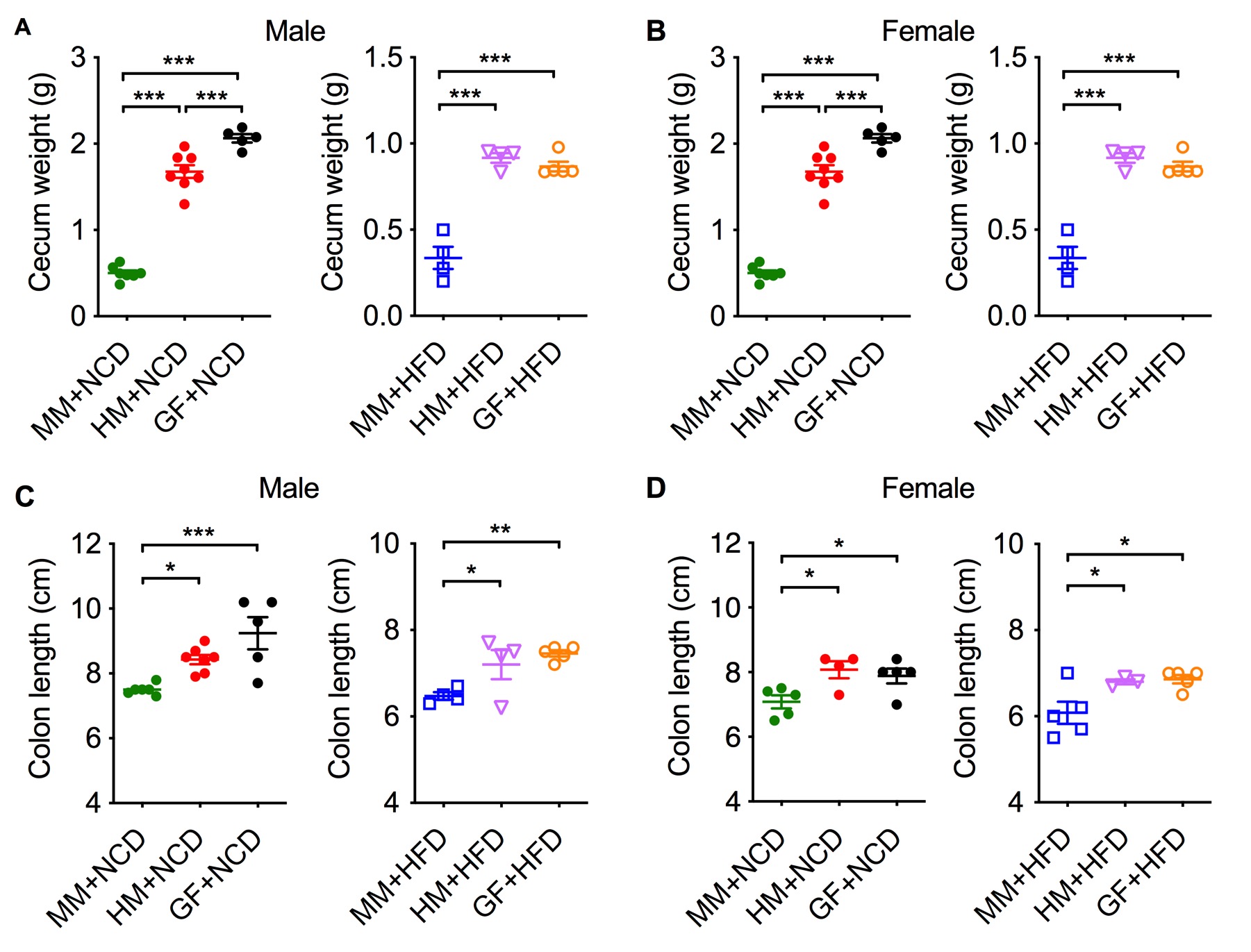

### Supplementary Figure 3

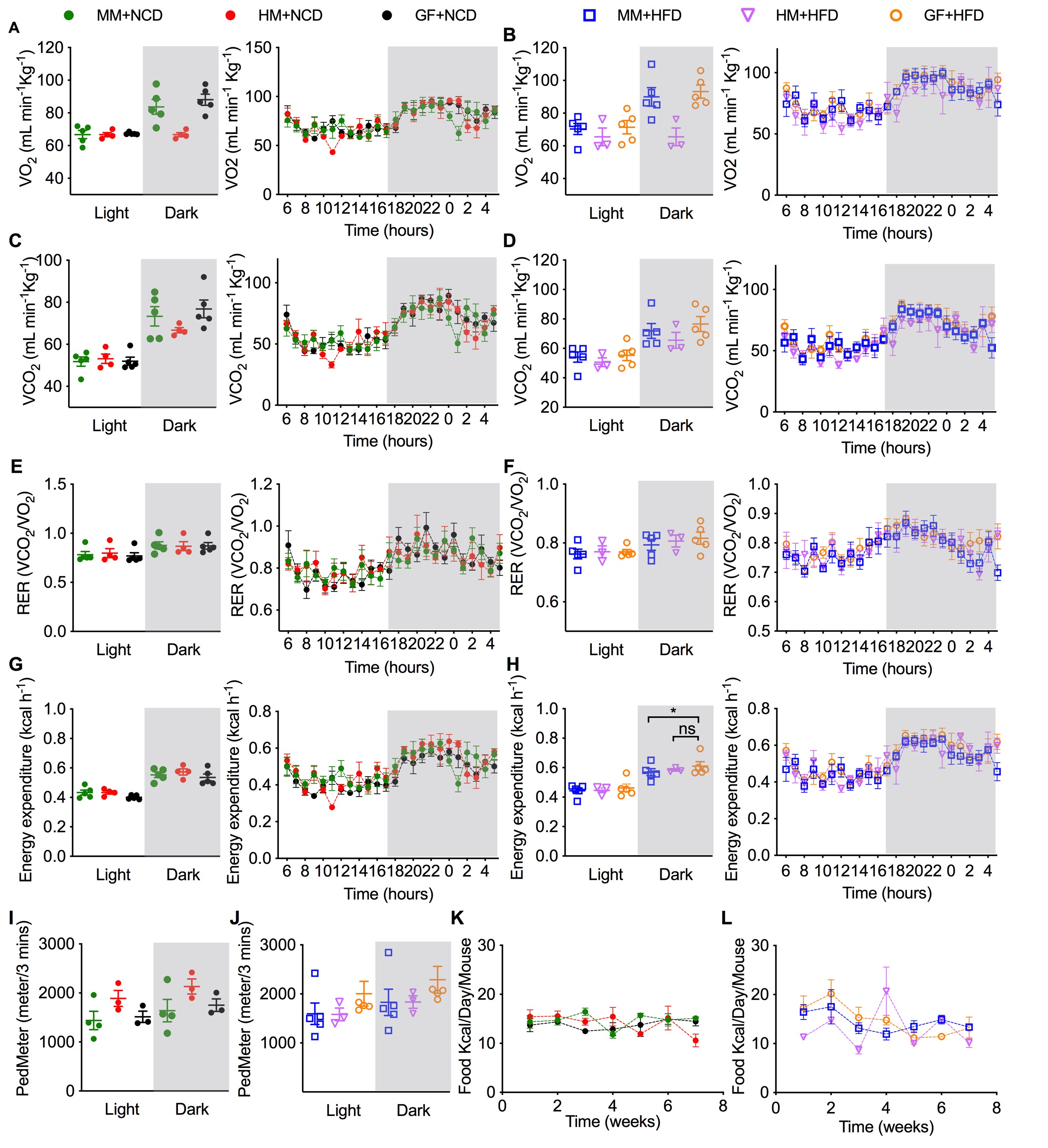

### Supplementary Figure 4

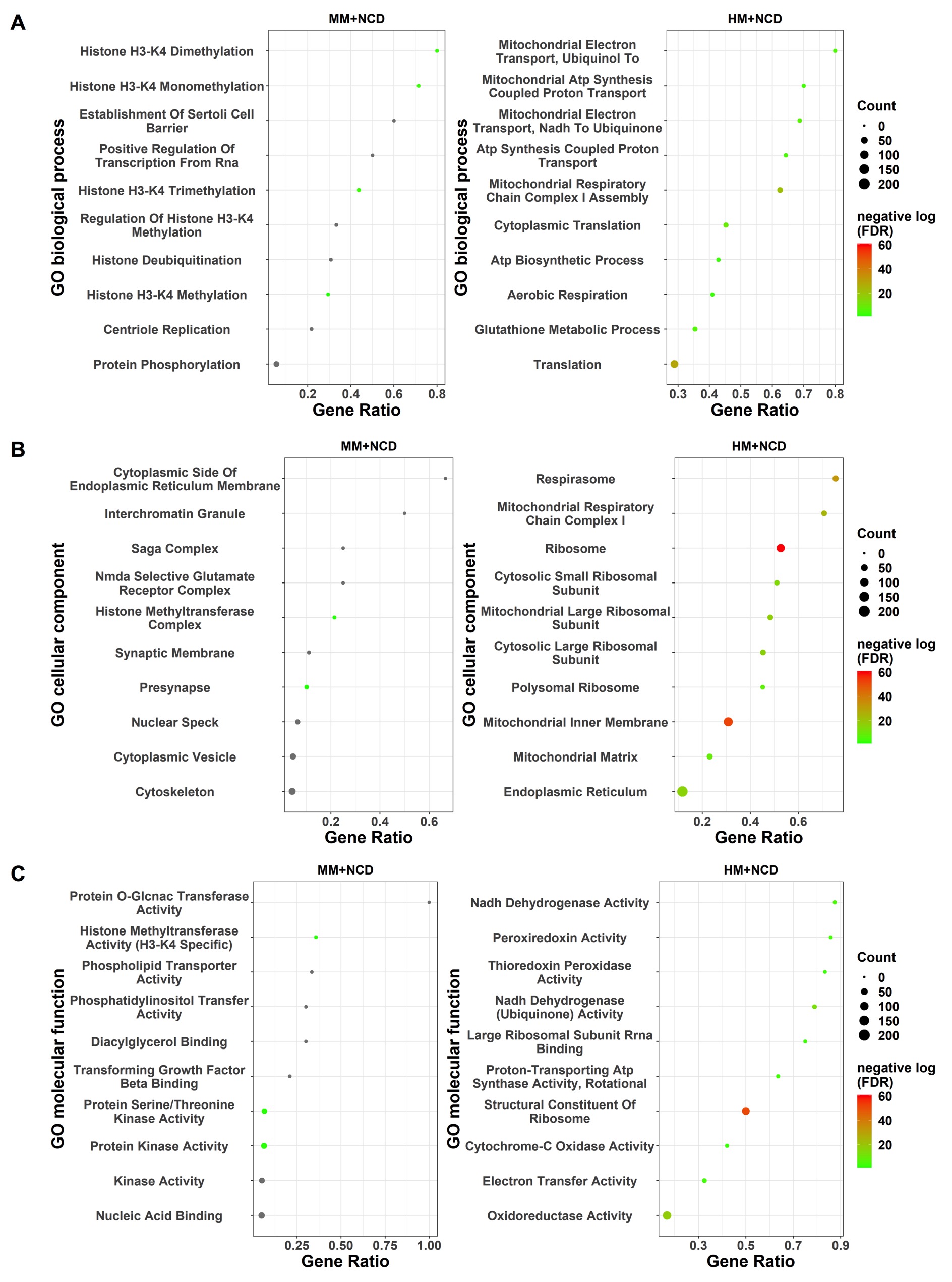

### Supplementary Figure 5

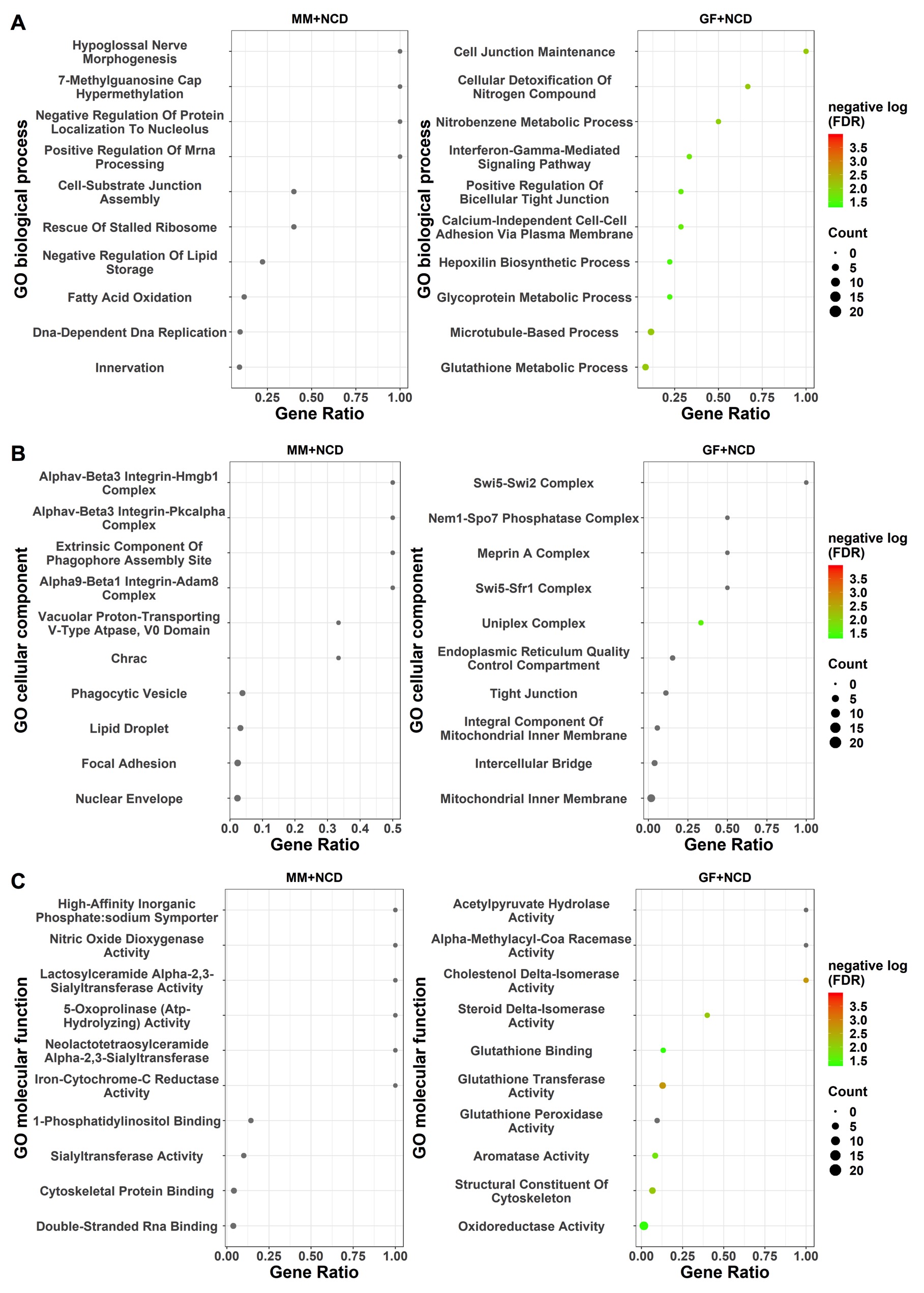

### Supplementary Figure 6

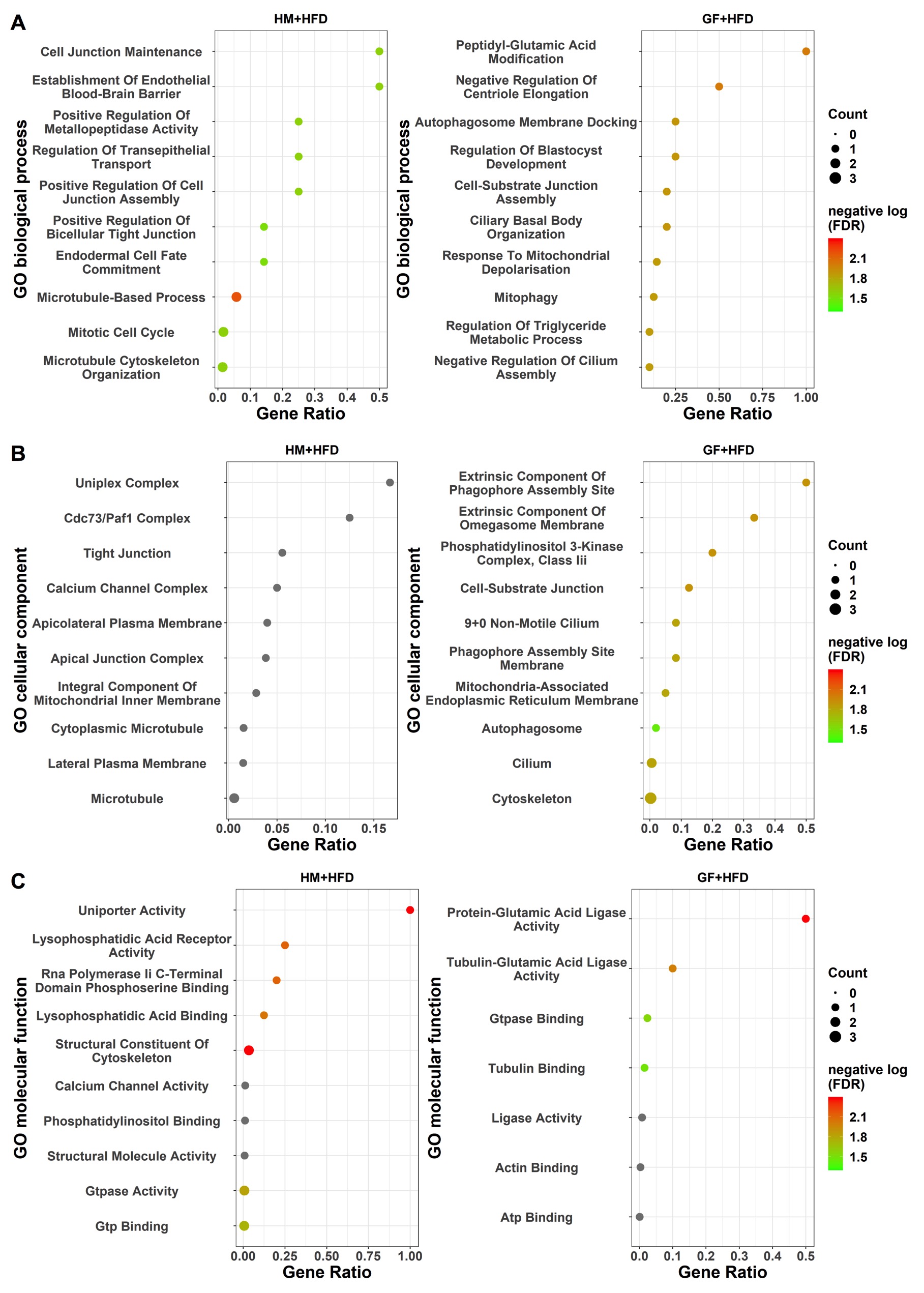
